# Asymmetric trichotomous data partitioning enables development of predictive machine learning models using limited siRNA efficacy datasets

**DOI:** 10.1101/2022.07.08.499317

**Authors:** Kathryn R. Monopoli, Dmitry Korkin, Anastasia Khvorova

## Abstract

Chemically modified small interfering RNAs (siRNAs) are promising therapeutics guiding sequence-specific silencing of disease genes. However, identifying chemically modified siRNA sequences that effectively silence target genes is a challenge. Such determinations necessitate computational algorithms. Machine Learning (ML) is a powerful predictive approach for tackling biological problems, but typically requires datasets significantly larger than most available siRNA datasets. Here, we describe a framework for applying ML to a small dataset (356 modified sequences) for siRNA efficacy prediction. To overcome noise and biological limitations in siRNA datasets, we apply a trichotomous (using two thresholds) partitioning approach, producing several combinations of classification threshold pairs. We then test the effects of different thresholds on random forest (RF) ML model performance using a novel evaluation metric accounting for class imbalances. We identify thresholds yielding a model with high predictive power outperforming a simple linear classification model generated from the same data. Using a novel method to extract model features, we observe target site base preferences consistent with current understanding of the siRNA-mediated silencing mechanism, with RF providing higher resolution than the linear model. This framework applies to any classification challenge involving small biological datasets, providing an opportunity to develop high-performing design algorithms for oligonucleotide therapies.

## INTRODUCTION

Small interfering RNA (siRNA) drugs guide potent and specific silencing of disease-related genes. siRNAs direct gene silencing by loading into the RNA-induced silencing complex (RISC) and binding the target site of an mRNA via complementary base pairing (1–3) (Figure 1). RISC then cleaves the target transcript, triggering mRNA degradation. With the recent FDA approval of four siRNA drugs (patisiran, givosiran, lumasiran, inclisiran) for liver-associated indications, and many other siRNAs in late-stage clinical trials, siRNAs have become one of the most promising drug modalities.

**Figure 1.**
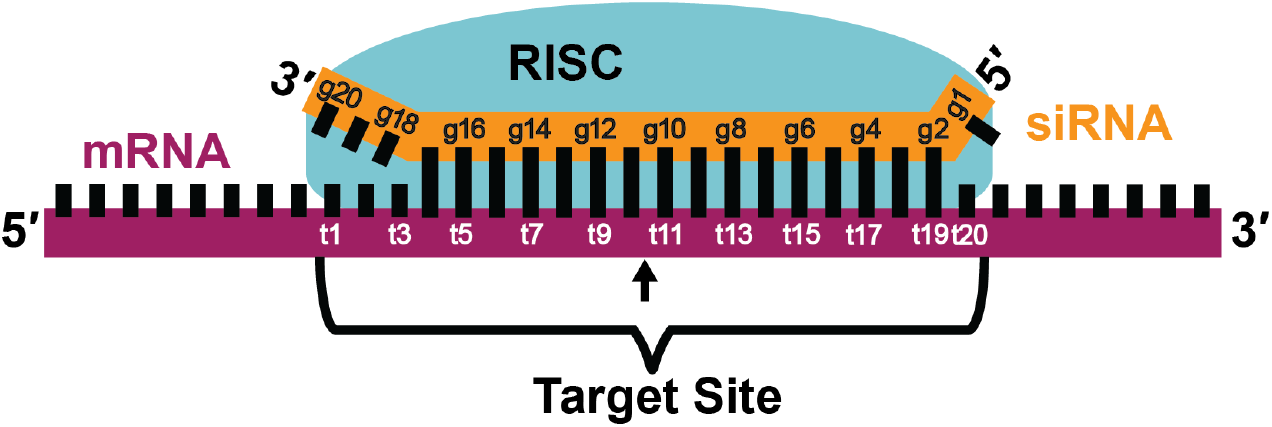
siRNA sequence data used in models and scoring scheme. A 20nt siRNA, when incorporated into the RNA-induced silencing complex (RISC), binds target mRNA via complementary base pairing between positions t2 and t17. The 20nt Target Site used for training siRNA design models is indicated. The siRNA guide strand positions are numbered (g1-g20). The mRNA target positions are numbered (t1-t20). The sequence of this region is used to train all models. Arrow indicates location of mRNA cleavage by the RISC between positions t10 and t11.

Despite siRNA sequence being a predictor of efficacy *(i*.*e*., degree of target gene silencing) (4), identification of siRNA target sites that effectively reduce target gene expression *in vivo* remains a key bottleneck in siRNA drug development. Therapeutic siRNAs must be fully chemically modified to increase stability and bioavailability *in vivo* (5–7), and tolerability of chemical modifications is a primary factor limiting siRNA efficacy. Although publicly available algorithm can accurately determine effective native siRNAs, their prediction accuracy is poor when applied to chemically modified compounds (4). Moreover, early “first generation” siRNA design algorithms often employed simple learning architectures such as linear models, which cannot describe complex sequence relationships that might underlie siRNA efficacy (8–13).

Machine learning (ML) approaches that leverage non-linear models are highly powerful in fitting complex patterns in the data (14–18), and can be applied to build algorithms of superior predictive power. Supervised ML methods focused on classification (*e*.*g*., effective versus ineffective) achieve this by training a model on a labeled dataset, with each data point assigned to a specific class label. Yet, many supervised machine learning methods require hundreds or thousands of labeled data points to build a model that accurately classifies the previously unlabeled data; and thus, are difficult to apply to modified siRNA efficacy datasets. Most data are industry generated and thus proprietary with the cost of synthesis and screening further limiting data availability to the research community. Studies applying ML to non-modified siRNA data have increased dataset size by combining heterogenous siRNA data (generated from structurally and chemically different compounds using different assays/conditions) (19–22). However, applying ML to heterogeneous data can generate noise, which reduces prediction accuracy.

Here, we apply a supervised ML approach to a small chemically modified siRNA efficacy dataset (n=356) by using the data itself to inform the classification process. Introducing a two-threshold (or trichotomous) model combined with a systematic assessment of a range of classification thresholds overcomes bias introduced by defining a single *ad-hoc* threshold. This trichotomous scheme enables selection of the optimal threshold pair to significantly reduce noise generated from using small siRNA datasets where large variability in signal is common. The resulting ML model showed high predictive power and outperformed a linear classification model built from the same data. To assess model validity and understand biological mechanisms underlying model results, we evaluated features dictating model prediction using a novel method for extracting sequence position-base preferences. This feature extraction method employs an evaluation-centered protocol that is agnostic to model type in contrast to the previous approaches. The two-threshold framework presented here can be applied to any noisy biological dataset to build powerful ML models and is designed to perform well even on small datasets. Such a framework unlocks the fully utility of siRNA datasets that are typically small and noisy to better understand siRNA mechanisms and design next-generation nucleic acid therapeutics.

## MATERIAL AND METHODS

### Dataset Acquisition

siRNA sequences and corresponding efficacy data used in this analysis were obtained from a publicly available dataset (4). These siRNAs were evaluated for their efficacy in target gene silencing in HeLa cells using a dual luciferase assay as described previously (4).

### Efficacy Threshold selection

Pairs of thresholds (effective – h_1_ – and ineffective – h_2_) were selected such that they were evenly distributed from the lowest reporter expression value in the dataset (4%) to the highest (120%) to maintain approximately the same number of points within each group (23-24). Nine h_1_ thresholds were defined to include siRNAs with report expression values: ≤15%, ≤22%, ≤29%, ≤35%, ≤40%, ≤46%, ≤53%, ≤65%, ≤82%. The nine h_2_ thresholds were defined with the same distribution and included all siRNAs with reporter expression values: >15%, >22%, >29%, >35%, >40%,>46%, >53%, >65%, >82%.

In the trichotomous partitioning scheme presented in this manuscript, siRNAs with reporter expression values greater than or equal to the effective threshold but less than or equal to the ineffective threshold were classified as undefined. All permutations of non-overlapping threshold pairs were considered (*i*.*e*., h_1_ ≤15%, h_2_ >15% was considered but h_1_ ≤35% and h_2_ >22% was not).

### Feature Parameterization

For each siRNA sequence, the 20-nt target site of the target mRNA was extracted (Figure 1) to generate feature vectors for training the model. Specifically, the sequences were encoded into basic binary features using the following protocol (25). A’s were represented as [1,0,0,0], U’s were represented as [0,1,0,0], C’s were represented as [0,0,1,0], and G’s were represented as [0,0,0,1]. For each sequence, arrays of encoded bases were appended in the order they appear in the sequence to form the final 80-dimensional feature vector, representing the full 20 nt target site sequence. Feature vectors where labeled with previously described activity classifications (effective/ineffective/undefined).

### Assessment Protocol

A training set containing 75% of the data (267 siRNAs) and holdout dataset containing 25% (89 siRNAs) were randomly selected using the Scikit-Learn *train_test_split* method, which ensured unbiased random partitioning of data into desired proportions and enabled equal distribution of effective and ineffective siRNAs when changing both h_1_ and h_2_ thresholds.

The training dataset was used in K-fold cross-validation during which it was partitioned randomly into 10 K groups using the Scikit-Learn *KFold* method, which ensured random and even partitioning of the data. To ensure all 267 siRNAs from the training set were included in cross-validation, six groups contained 30 siRNAs and three groups contained 29 siRNAs. During partitioning of K groups, dataset classification was considered to ensure test groups were balanced with approximately equal numbers of effective and ineffective siRNAs.

Model performance was assessed by <u>A</u>rea <u>U</u>nder the <u>P</u>recision-<u>R</u>ecall <u>C</u>urve adjusted (AUCPR_adj_) measure. To compute AUCPR_adj_, the precision-recall curves were plotted using the *Matplotlib* package (26), followed by computing the AUCPR values using the *auc* function from Scikit-Learn (27). Values of AUCPR_adj_ were normalized to reflect the range from 0 to 100. The color scheme was designed to follow the same range from 0 (blue) to 100 (yellow).

### Machine Learning Model Training

Using the feature vectors, the supervised learning models were trained using the *Random Forest Classifier* from the Scikit-Learn Python Package (27). All RF models were trained with the following default parameters: 200 total trees with a maximum tree depth of 3 nodes, and at least one sample per leaf. Model performance was also evaluated solely on the training set during training (Supplementary Figures S8 and S9). The linear classifier models were trained using a published method that leverages an *ad-hoc* function and the three activity classification groups (4). This linear method was selected because it was previously applied to the siRNA dataset used here (4).

### Position-base preference determination

The direct feature extraction method applied previously to a linear model (4) is not applicable in the case of more advanced non-linear RF model, therefore we developed an alternative “proxy” feature extraction method that is agnostic to the model type (see Results). When applied to the linear model, the new feature extraction method shows comparable performance determining the same important features (Supplementary Figure S6). Feature importance from both linear and RF models – regardless of derivation method – were normalized to values between -100 and 100. Feature importance between -20 and 20 for all models were set to zero to minimize noise from low-weight features.

## RESULTS

### siRNA efficacy dataset and two-threshold class annotation

A chemically modified siRNA efficacy dataset consisting of 356 target sequences (4) was used for ML model training. The dataset comprises compounds targeting 17 genes, with an average of 15 siRNAs per gene. Sequences were designed with minimal constraints, mostly limited to favoring low GC content. All siRNAs were designed with the same chemical modification pattern. The 20-nt target site sequence for each siRNA was used as a training set for the supervised machine learning model. Base preferences at each position of a target site (Figure 1) were used as features—4 bases × 20 nt positions = 80 position-base features—to encode representation of each data point (see Methods for feature parametrization) (4, 8).

siRNA efficacy was determined by a dual luciferase reporter assay (4), and defined as reporter expression in cells treated with siRNA as a percent of reporter expression in untreated cells. siRNA efficacies ranged from 4% to 120% reporter expression, with a mean and median of 44% and 40%, respectively (Figure 2A). The luciferase reporter allows unification of the siRNA dataset by using a single experimental measure of efficacy. The average percent error was 3%, which is common for this type of data. Nevertheless, the dataset was relatively noisy, with individual siRNA efficacy values varying up to 16% (Figure 2A).

**Figure 2.**
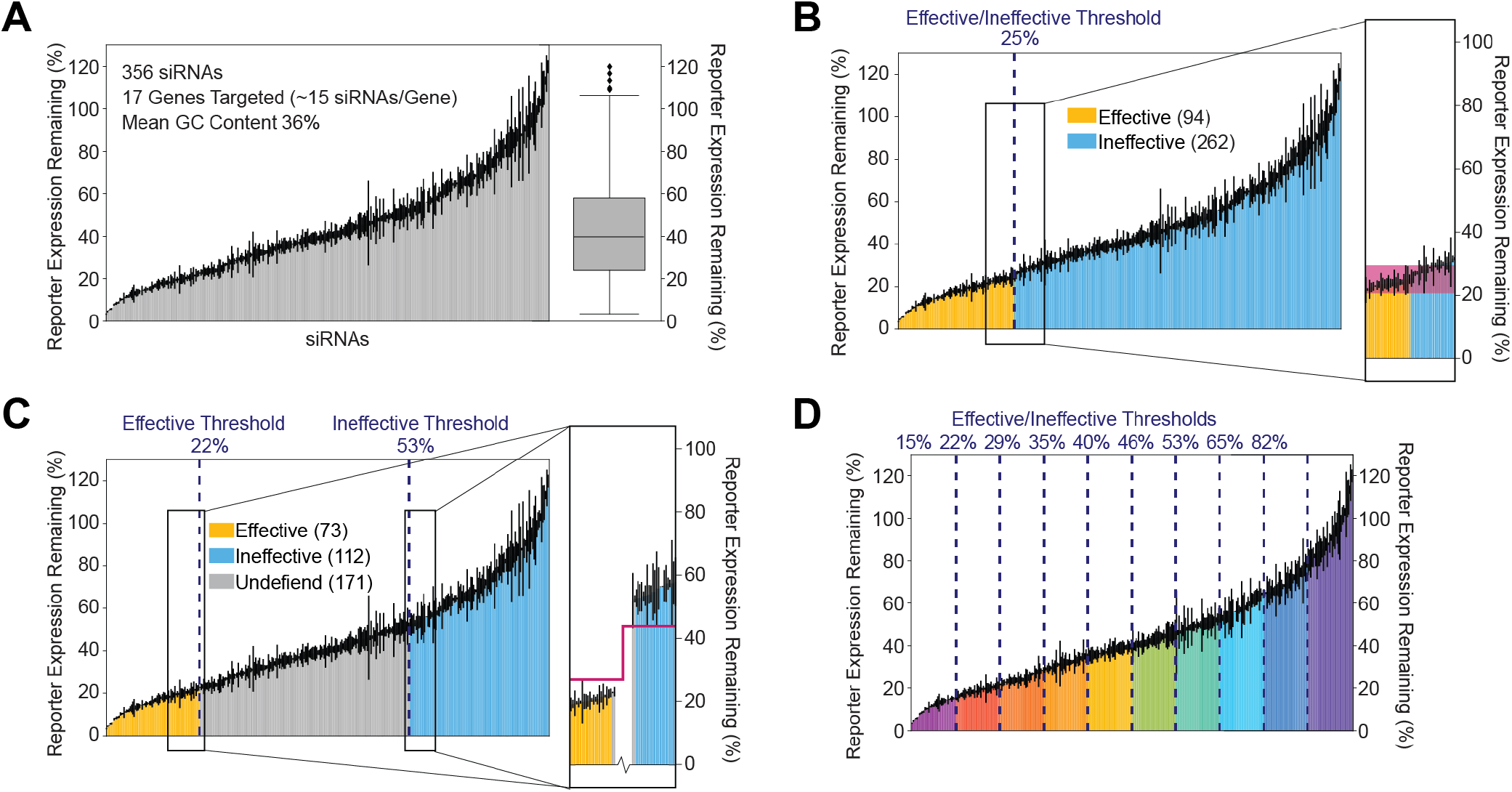
Noisy data and intermediate values challenge siRNA classification. (A) Gene silencing efficacy for 356 chemically modified siRNAs evaluated targeting 17 different genes (∼15 siRNAs/gene) in HeLa cells using a dual luciferase assay normalized to nontreated cells. Each bar represents efficacy of a single siRNA sequence averaged over three independent measurements with error bars depicting the standard deviation. Box and whisker plot depicts distribution of siRNA efficacies across the entire dataset. (B, C) Data in panel A classified as effective (yellow), ineffective (blue), or undefined (grey) by the thresholds indicated (blue dotted lines; threshold reporter expression % indicated at top). Number of siRNAs in each class indicated in parentheses. Inset shows regions around thresholds in greater detail. Shaded maroon boxes indicate regions with overlapping noise in the effective and ineffective classes (from maximal standard deviation value in the effective class to minimal standard deviation value in the ineffective class). Maroon bars indicate regions without overlap between effective and ineffective classes. (D) Data in panel A with all nine evenly spaced thresholds used in evaluation (blue dotted lines; threshold reporter expression remaining % indicated at top). Spans between thresholds defined 35-36 siRNAs. Threshold pairs contained all possible combinations of nonoverlapping effective/ineffective thresholds, resulting in 45 possible combinations. effective classes contained all siRNA sequences less than or equal to the threshold. Ineffective classes contained all siRNA sequences greater than the threshold.

Consistent with other siRNA efficacy datasets, distribution of the data did not provide a clear threshold for classification (Figure 2A). A biologically reasonable threshold of 25% reporter expression (Figure 2B) would define 94 effective siRNAs. However, data points at both sides of the threshold have large overlap in error bars (inset, Figure 2B). This noise makes data points around the threshold indistinguishable – a point directly to the right of the threshold is no different from a point to the left of it. Thus, this classification will result in a subset of sequences with biologically equivalent efficacies distributed to different classes.

To overcome this issue, we applied a non-conventional trichotomous grouping method that uses two independently selected thresholds, h_1_ < h_2_. In this approach, one threshold defines effective siRNAs (h_1_), and another threshold defines ineffective siRNAs (h_2_). All siRNAs lying between these thresholds are classified as “undefined” and excluded from model development (Figure 2C). This results in two clearly distinct groups with no “noise overlap” (inset, Figure 2C). To determine optimal effective and ineffective siRNA threshold values, different pairs of h_1_ and h_2_ efficacy thresholds (from 15% to 82% reporter expression) were selected for testing (Figure 2D). siRNAs with reporter expression values less than the selected h_1_ threshold were labelled ‘effective’, while those with values greater than the selected h_2_ threshold were labeled ‘ineffective’. All permutations of effective and ineffective threshold pairs were systematically evaluated, excluding threshold combinations that would classify the same siRNA(s) into both effective and ineffective classes (*e*.*g*., h_1_ ≤15%, h_2_ >15% was considered but h_1_ ≤35% and h_2_ >22% was not). Thus, a total of forty-five threshold combinations were considered. The size of the dataset used for model building was affected by threshold selection: models built with the most stringent threshold (h_1_ ≤15% and h_2_ > 82% – hereon written as 15/82) were built using data from the fewest siRNAs, while models built using identical effective and ineffective thresholds use the whole dataset (*e*.*g*., 15/15, 22/22, etc.).

### Pipeline for classification model development

We evaluated the impact of all 45 threshold combinations on model performance using the pipeline in Figure 3. For each threshold pair (Figure 3A), a supervised ML model employing random forest (RF) classification was built. RF was selected because this ML model type is known to achieve a learning plateau in the fastest way, requiring the fewest number of training examples among all nonlinear ML methods (28). RF models use decision trees to partition data by their features to classify the data. In our analysis, the RF models partition data by target site position-base features to classify siRNAs as positive (*i*.*e*., effective) or negative (*i*.*e*., ineffective). RF performs well on data with a large number of features, and the trees and branching structure of RF have the potential to capture complex interactions (*e*.*g*., sequence motifs) that a simpler linear model cannot (18).

**Figure 3.**
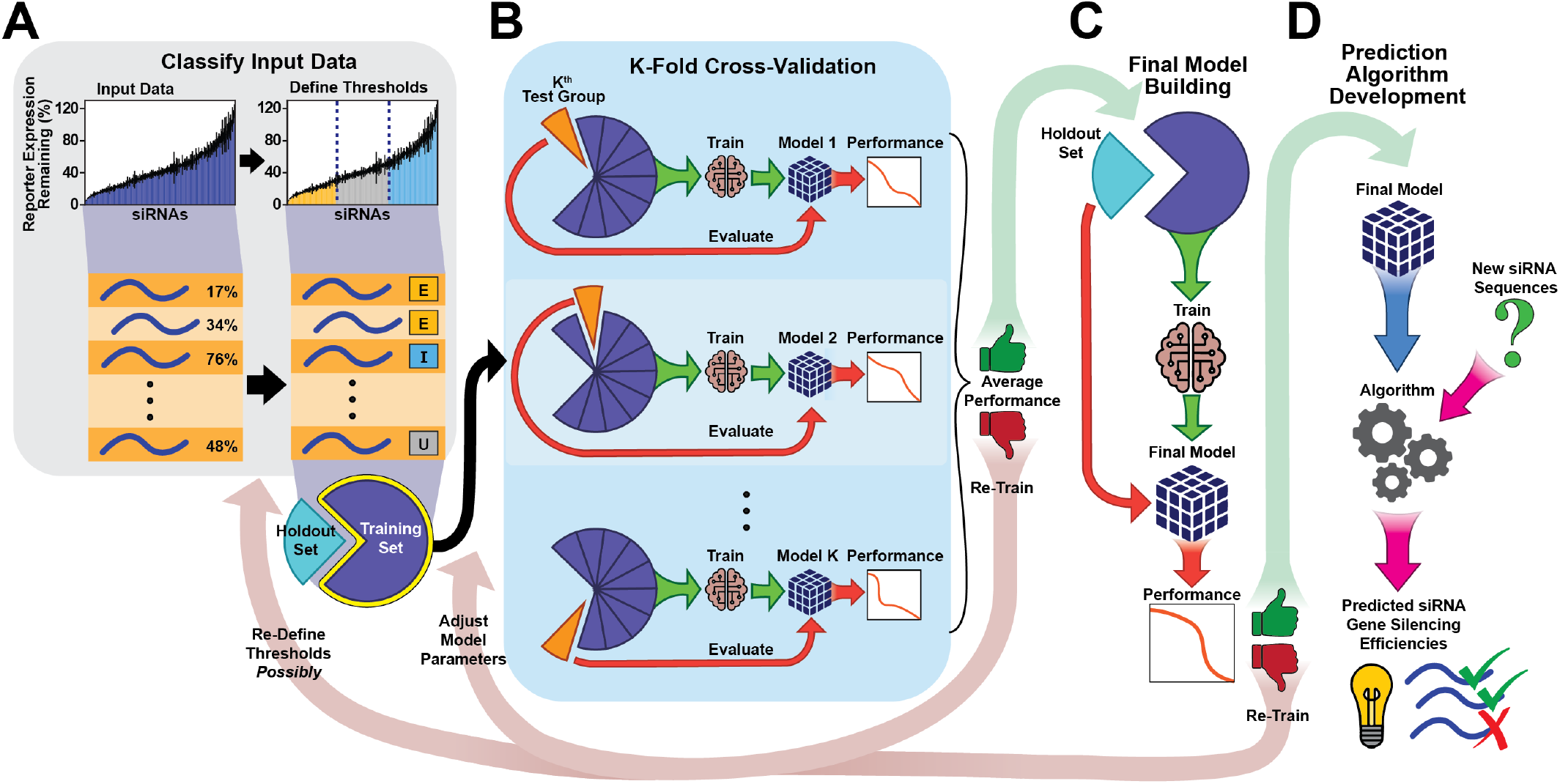
Schema for training machine learning model to produce efficacy prediction algorithm for a single threshold combination. (A) Input siRNA sequences with experimentally determined gene silencing efficacy are classified using predefined thresholds (dotted lines) into three groups: effective (E), ineffective (I), and undefined (U). Classified data are partitioned randomly into holdout (25% of data) and training (75% of data) sets. (B) Training data are split into K (10) groups of equal size. K^th^ test group consisting of 10% (one group) of the training data is held out (orange pie slice) and a model is trained on the remaining 90% (nine groups) of the training data. Model performance is evaluated using the K^th^ test group. If average performance of all K models is acceptable final model building can proceed, otherwise the model must be re-trained. (C) Using the full training set, the final model is built, and its performance is evaluated on the holdout set. If the performance is acceptable prediction algorithm development can proceed using the model, otherwise the model must be re-trained. (D) The final model is used to build an algorithm to predict gene silencing efficacies of siRNA sequences whose efficacy has not been experimentally determined.

For model development and independent validation, the dataset was partitioned into a training set and holdout set, consisting of 75% and 25% of the data, respectively (Figure 3A). The data were split to ensure equal distribution of effective and ineffective siRNAs (per selected threshold pair for that model) into the training and holdout sets to minimize biases and optimize model development (see Methods for assessment protocol).

Because the training and holdout sets inherently have different characteristics, bias can be introduced into the model during partitioning. This is particularly true for small, diverse datasets, as anomalies existing in only a few data points (as few as 5 siRNAs) can cause a model to underperform. This bias is minimized using K-fold cross-validation – an iterative process in which the training set is randomly partitioned into K groups of equal size, then K rounds of model building are performed using K-1 groups in training and 1 group in testing (29). The testing group is then used for model evaluation (Figure 3B). For siRNA prediction models, K was set to 10, a typical number for a dataset of this size, and the cross-validation process was repeated a total of 10 times such that each partition served as the testing group once. The average performance of the K models was then analyzed (see section on novel scoring metric for model evaluation). Default values of the standard parameters of a RF model, depth of the tree and the number of trees, were chosen because altering these parameters did not impact model performance (data not shown). Following K-fold cross-validation, final model training was performed with the entire training set evaluated on the holdout set (Figure 3C). Because the holdout set was not involved in K-fold cross-validation, model performance on this set is a strong indicator of model generalization (*i*.*e*., performance on future unseen data). The final model is then used to build an algorithm to predict effective and ineffective siRNAs (Figure 3D).

### A novel scoring metric, AUCPR_adj,_ for model evaluation across two-threshold combinations

Accuracy, a popular model performance metric that measures correct versus total predictions, is misleading in the context of large class imbalances. This is of particular concern for fully modified siRNA efficacy datasets, like the one used here (33), in which there are many more ineffective siRNAs than effective siRNAs for a target transcript (34, 35). Another popular model performance metric is area under the Receiver Operating Characteristic (ROC) curve, which plots the true positive rate (also called recall) against the false positive rate (36) Yet, like accuracy, ROC curves do not account for imbalanced data, overestimating model performance in datasets dominated by positively classified values (*i*.*e*., permissive efficacy threshold).

Precision-Recall (PR) curve is a better metric for siRNA design models (38). *Recall* (Equation 1) depicts the fraction of siRNAs correctly predicted as effective with respect to all effective siRNAs in the dataset (37) A model producing a large number of false negatives (effective siRNAs classified as ineffective) will have low recall. *Precision* (Equation 2) is the fraction of siRNAs correctly predicted as effective with respect to all siRNAs predicted to be effective (37). A model producing a large number of false positives (ineffective siRNA classified as effective) will have low precision. When combined, recall and precision consider both false negatives and false positives to capture both class types, overcoming class imbalance issues in model evaluation.

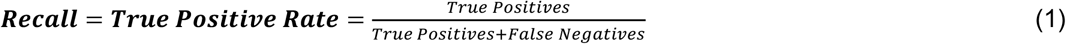

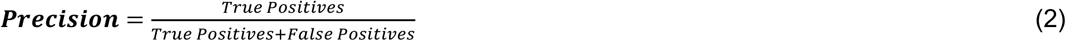

The goal of siRNA design is to identify a panel of siRNA sequences that effectively silence a target gene. A strong model for this use need not identify *all* possible effective siRNA sequences, but the sequences it does identify should have a high probability of being effective. Such a model will prioritize high precision (majority of siRNA classified as effective are effective), over high recall (identifying all possible effective siRNA).

The area under the precision-recall curve (AUCPR) converts the PR curve to a single numeric value (38). Higher AUCPR usually indicates a better performing model (Figure 4A). Unfortunately, in the context of vastly different h_1_ thresholds, the AUCPR of different curves are not comparable because changing h_1_ affects the precision when recall equals 1 (P_R=1_), causing more permissive h_1_ thresholds to automatically generate a higher AUCPR (blue vs gold curves, Figure 4B), and allowing two models with different thresholds and performance to potentially produce identical AUCPR (Figure 4C). To overcome this, we adjusted AUCPR to the P_R=1_ by subtracting the area defined by precision at maximum recall, creating a metric called AUCPR_adj_ (Figure 4D, Equation 3). This normalization maintains proper performance assessment of models built with the same h_1_ threshold (Figure 4E), corrects for poor (low AUCPR_adj_) assessment of overfit and underfit models (Figure 4F), and distinguishes between models with otherwise identical AUCPRs (Figure 4G).

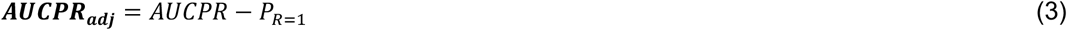

To simplify comparison between different models, we further normalize AUCPR_adj_ to 1:100 scale, creating a standalone performance metric (color scale bar, Figure 4D). In addition, contingency tables, which quantify true positive, false positive, true negative, and false negative groups, are used to define the source of poor performance (Supplementary Figures S1 and S2).

**Figure 4.**
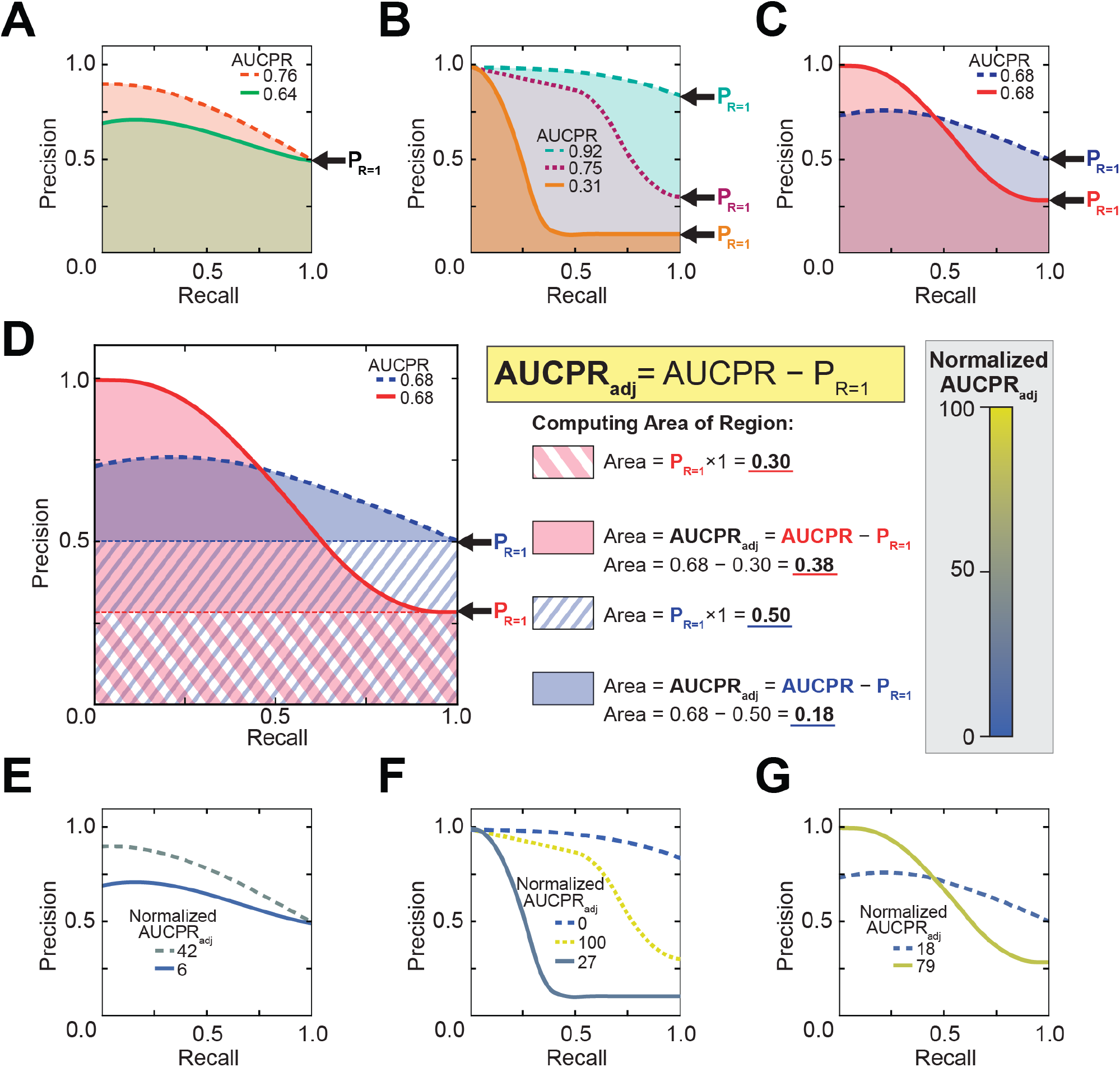
Adjusted area under the precision-recall curve overcomes challenges of model evaluation using precision-recall curves. (A) Precision-recall curves of two models developed using the same threshold values. Curves depict model performance for a better performing model (orange dashed curve) and a worse performing model (green curve). Areas under the precision-recall curve (AUCPR) represented by shaded regions are indicated at top. Arrow identifies precision when recall equals one (P_R=1_). (B) Precision-recall curves of an overfit model with AUCPR overestimating performance (teal dashed curve), an underfit model with AUCPR underestimating performance (gold line), and a top-performing model (purple dashed curve). Arrows identify P_R=1_ values for the curve of the corresponding color. (C) Precision-recall curves for models developed with different effective thresholds: a more stringent threshold (red curve) and a more permissive threshold (blue dashed curve) resulting in identical AUCPRs. (D) Same data as in (C) depicting AUCPR_adj_ (red and blue shaded regions) derivation by subtracting the area defined by the P_R=1_ (red/white and blue/white striped regions) from the corresponding curve’s (red or blue) AUCPR. General formula for computing AUCPR_adj_ is described (yellow box). Detailed AUCPR_adj_ derivations for the blue and red curves provided (middle). Color bar represents the scheme used throughout this manuscript for color-coding curves by AUCPR_adj_ values normalized between 0 and 100. (E, F, G) Same curves as in A, B, C, respectively, colored by normalized AUCPR_adj_.

### Performance of RF model for siRNA prediction is highly affected by classification thresholds

Using AUCPR_adj_, we found averaged K-fold cross-validation model performance (Figure 5 and Supplementary Table S1) and final model performance on the holdout dataset (Figure 6 and Supplementary Table S2) to be generally similar. There was an overall trend of higher performance for models built with the most stringent threshold pairs (small h_1_/large h_2_; top left curves, Figures 5 and 6), while models built with all other threshold combinations performed poorly (bottom left, top right, and center curves, Figures 5 and 6).

**Figure 5.**
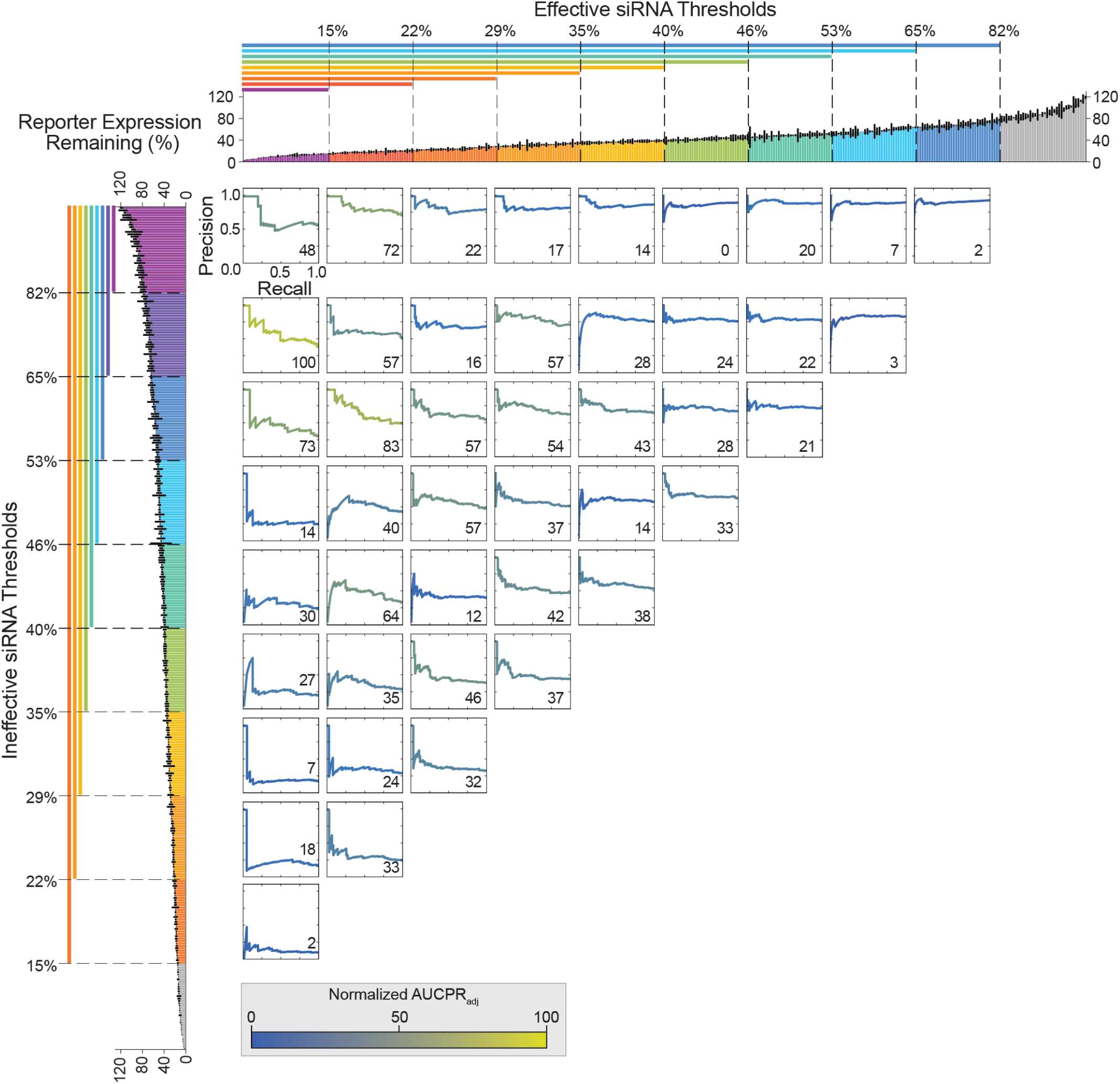
Model performance from K-fold cross-validation per classification threshold. Precision recall curves for random forest classifiers trained on 9/10^th^ training groups and evaluated on corresponding 1/10^th^ test groups averaged over 10 rounds of model building. Each curve represents performance of models trained using different effective and ineffective siRNA threshold pairs. Curves are colored by normalized AUCPR_adj_ (indicated at bottom right of each curve). Color bar depicts performance by normalized AUCPR_adj_: Values of AUCPR_adj_ were normalized to reflect the range from 0 to 100. The color scheme was designed to follow the same range from 0 (blue) to 100 (yellow). Bar plots at top and left depict all siRNA target expression data (as in Figure 2D) colored by effective (top) or ineffective (left) thresholds. Precision recall curves are aligned to these bar plots to indicate the effective and ineffective thresholds used for training of the corresponding curve’s model. Thresholds are inclusive of all data with expression values less than (for effective thresholds) or greater than (for ineffective thresholds) the threshold expression percentage. Data used to compute normalized AUCPR_adj_ values in Supplementary Table S1.

**Figure 6.**
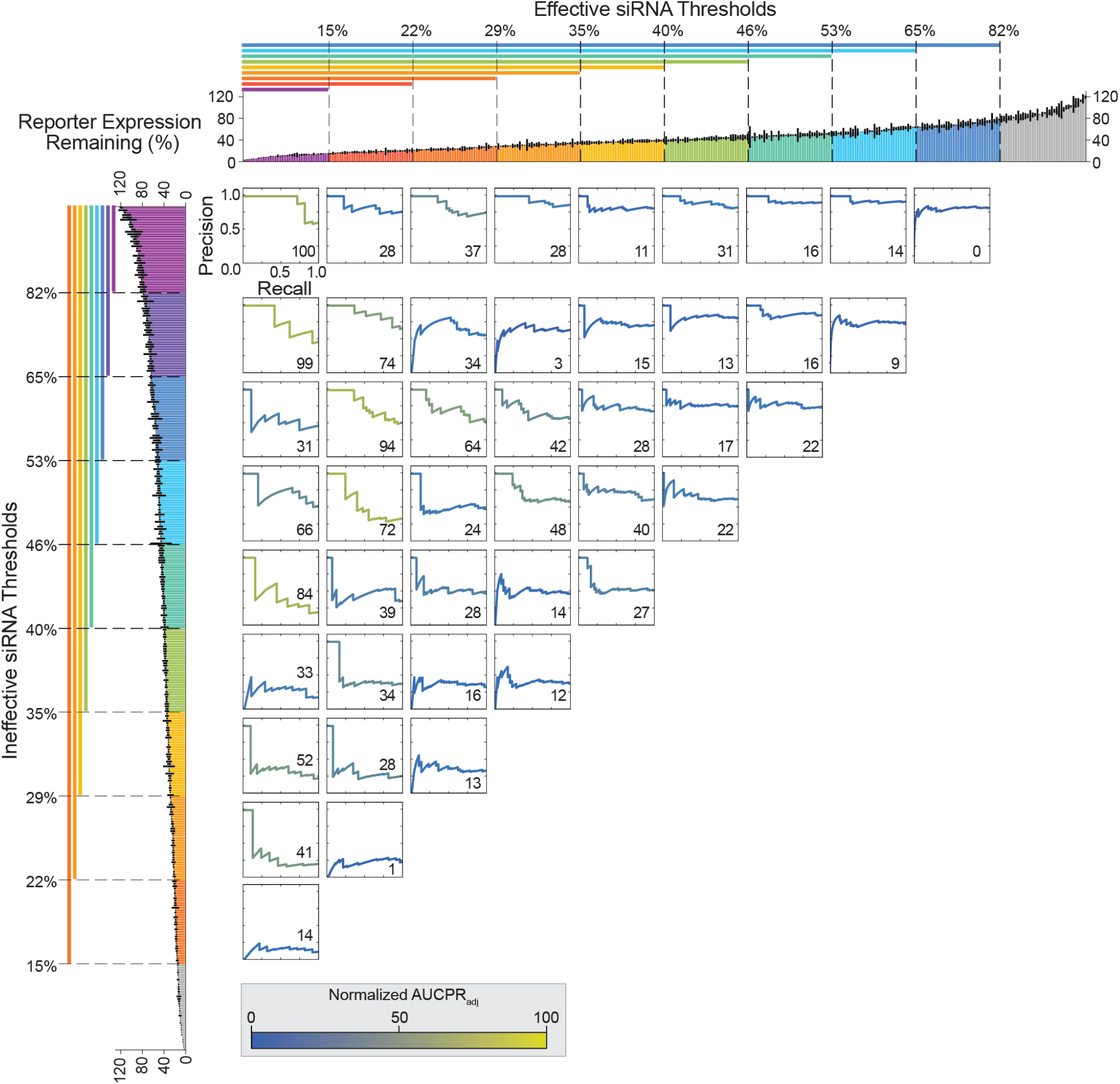
Model performance per classification threshold. Precision recall curves for random forest classifiers trained on entire training set and evaluated on holdout set. Each curve represents performance of a model trained using different effective and ineffective siRNA threshold pair. Curves are colored by normalized AUCPR_adj_ (indicated at bottom right of each curve). Color bar depicts performance by normalized AUCPR_adj_: Values of AUCPR_adj_ were normalized to reflect the range from 0 to 100. The color scheme was designed to follow the same range from 0 (blue) to 100 (yellow). Bar plots at top and left depict all siRNA target expression data (as in Figure 2D) colored by effective (top) or ineffective (left) thresholds. Precision recall curves are aligned to these bar plots to indicate the effective and ineffective thresholds used for training of the corresponding curve’s model. Thresholds are inclusive of all data with expression values less than (for effective thresholds) or greater than (for ineffective thresholds) the threshold expression percentage. Data used to compute normalized AUCPR_adj_ values in Supplementary Table S2.

Top-performing models – defined by a high AUCPR_adj_ – utilized threshold pairs that 1) reflect biologically reasonable definitions of siRNA efficacy (<30% reporter expression) and 2) exclude moderate-efficacy siRNA, which might be misclassified and introduce noise. The resulting models have the greatest power to distinguish effective and ineffective siRNAs, but come at the cost of excluding a larger amount of data from training. In K-fold cross-validation, the top performing threshold pairs (h_1_/h_2_) were 15/65, 15/53, 22/82, and 22/53 (Figure 5). The most stringent threshold pair, 15/82, did not perform well, likely due to the smaller dataset used. In final model evaluation, AUCPR_adj_ identified 15/82, 15/65, 15/40, 22/65, 22/53, and 22/46 as top-performing threshold pairs (Figure 6). The strong performance of the 15/40 threshold pair, which allows greater inclusion of moderate-efficacy siRNAs, was driven by the identification of true negatives (contingency table, Supplementary Figure S2). In fact, the model did not identify any true positives, suggesting the model would not likely perform well for effective siRNA identification. This exemplifies the challenges of model building with a limited dataset, where thresholding can further reduce the evaluation set size (to as few as 17 siRNAs in this assessment), and demonstrates that no evaluation metric is perfect. Considering multiple metrics – in this case AUCPR_adj_ and the contingency table – is critical for evaluating final model performance.

AUCPR_adj_ successfully identified three categories of poor-performing models. The first category of models utilized moderately effective and ineffective threshold pairs (center curves, Figures 5 and 6). Corresponding contingency tables (center tables, Supplementary Figures S1 and S2) show the models falsely classify many ineffective siRNAs as effective. This poor performance is likely due to the models including moderate-efficacy siRNAs. The two remaining categories were highly underfit or overfit models. Overfit models are identified by a high AUCPR with a high P_R=1_, and were built from threshold pairs producing a larger number of effective siRNAs (h_1_ = 53%, 65%, or 82%) (top right curves, Figures 5 and 6). This poor performance is likely due to overly permissive effective thresholds mislabeling ineffective siRNAs as effective. Underfit models are identified by their low AUCPR and P_R=1_, and were built from threshold pairs producing a larger number of ineffective siRNAs (h_2_ = 29%, 22%, or 15%) (bottom left curves, Figures 5 and 6).

Some threshold pairs produced models that performed notably worse on the holdout set than they did in cross-validation (22/65, 15/46, 22/46, and 15/40) (Figures 5 and 6). This is likely due to inherent differences between the holdout and training datasets that were not captured in the model during training, and reflects the small dataset size. These discrepancies are not due to unequal representation of effective and ineffective siRNAs in the holdout versus test groups in K-fold cross-validation (Supplementary Figures S1 and S2); equal representation of effective and ineffective siRNAs was maintained during partitioning of training and holdout sets as well as during k-fold partitioning (see Methods).

Evaluating model performance using ROC curves (and their corresponding AUC) did not provide a clear top performing model (AUC > 0.85) (36) (Figures S3 and S4), exemplifying the importance of selecting the proper metric for model evaluation. This is particularly striking when considering final model evaluations, where the most permissive ineffective siRNA thresholds showed some of the greatest AUCs (bottom left curves, Figure S4, Supplementary Figure S4) despite being underfit, as determined by their contingency tables.

### RF model outperforms linear classification model built from the same dataset

We next compared the performance of RF with a linear model to classify siRNA efficacy in this dataset (Figure 7). For this comparison, we selected a published linear classification method that leverages an *ad-hoc* function and utilizes the threshold pairs (4). We selected this linear method because it was previously applied to the siRNA dataset used here (4).

**Figure 7.**
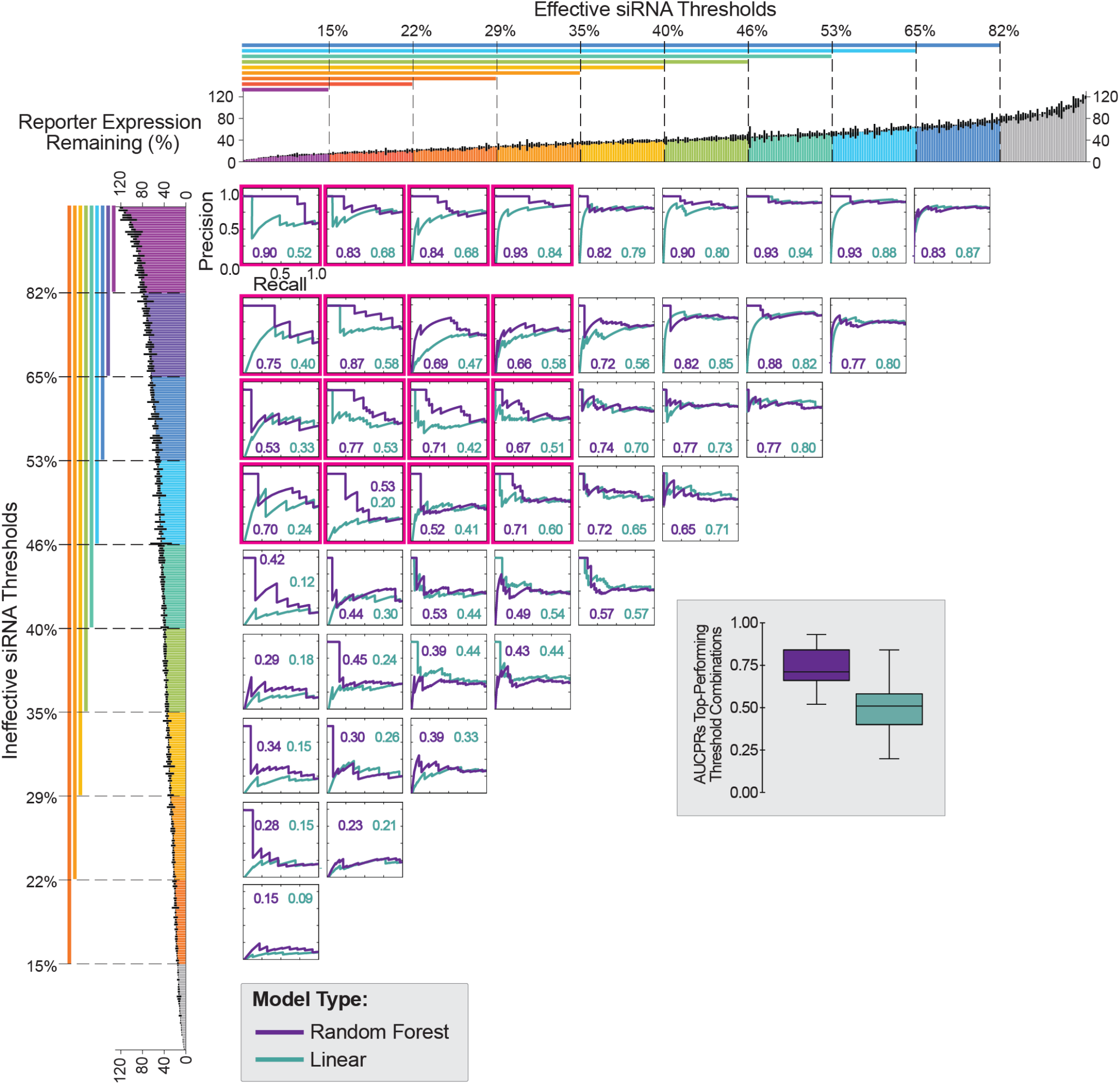
Comparing random forest and linear models. Precision recall curves for random forest classifiers (purple curves) and linear classifiers (teal curves) trained on entire training set and evaluated on holdout set. Each pair of over overlaid curves represent performances of models trained using a different effective and ineffective siRNA threshold pair. Bar plots at top and left depict all siRNA target expression data (as in Figure 2D) colored by effective (top) or ineffective (left) thresholds. Precision recall curves are aligned to these bar plots to indicate the effective and ineffective thresholds used for training of the corresponding curve’s model. Thresholds are inclusive of all data with expression values less than (for effective thresholds) or greater than (for ineffective thresholds) the threshold expression percentage. Areas under the precision recall curves (AUCPRs) are indicated in bottom right corner of each plot and color-coded by model type. Box plots on right depict the distribution of AUCPRs for models built with the most stringent thresholds (boxed in pink).

As with the RF model, there was a general trend in higher performance of linear models built with the most stringent threshold pairs (top left curves boxed in pink, Figure 7). Overall, RF performed better than linear, as determined by higher mean and median AUCPRs for top performing models (box plot, Figure 7). While overall performance of the linear model was significantly worse than RF, it showed some predictive power with the same top performing thresholds, indicating that elimination of moderate efficacy siRNAs from model development is beneficial for both simple linear models and more sophisticated ML models.

### Visualization of siRNA position-base preferences driving models by proxy feature extraction

To better understand how the top-performing RF model identifies effective and ineffective siRNAs, we looked at 20-nt target site position-base preferences. While deriving this type of matrix is trivial for a linear model, feature extraction from any ML model is complex and model-dependent (39). To overcome this challenge, we devised a method to estimate preferences that provide a proxy for relative importance of different feature contributions in decision trees.

To start, feature vectors for each siRNA used in model development were obtained (see Methods). Next, the model was used to predict efficacy of each siRNA. From the known siRNA efficacy and selected h_1_ threshold, each siRNA prediction is placed into one of four classification groups (true positive, true negative, false positive, or false negative). Each of the four classification groups is then considered individually, and the feature vectors of the siRNAs in each group are averaged. The averaged weights of the vectors representing the false positives and false negatives groups are multiplied by -1 as these indicate incorrect predictions. The resulting vectors from the four classification groups are then summed, and the summed vector is transposed back to represent base frequencies at each position in the 20-nt target region (Supplementary Figure S5). The resulting base preferences were plotted as a matrix with positive weights indicating preference for a base at a particular position in effective siRNA, negative weights indicating disfavoring of that base, and weights with zero values indicating no preference for that base at the given position (Figure 8). This method relies on the output of the classification model, and is agnostic to the model type. Therefore, it can be applied to any classification model to extract feature preferences. To test the veracity of this proxy method, we compared the base preference matrix generated from a linear model by proxy and by direct extraction methods. The two matrices appear similar, indicating that the proxy method provides a reasonable approximation of features driving the model performance (Supplementary Figure S6).

**Figure 8.**
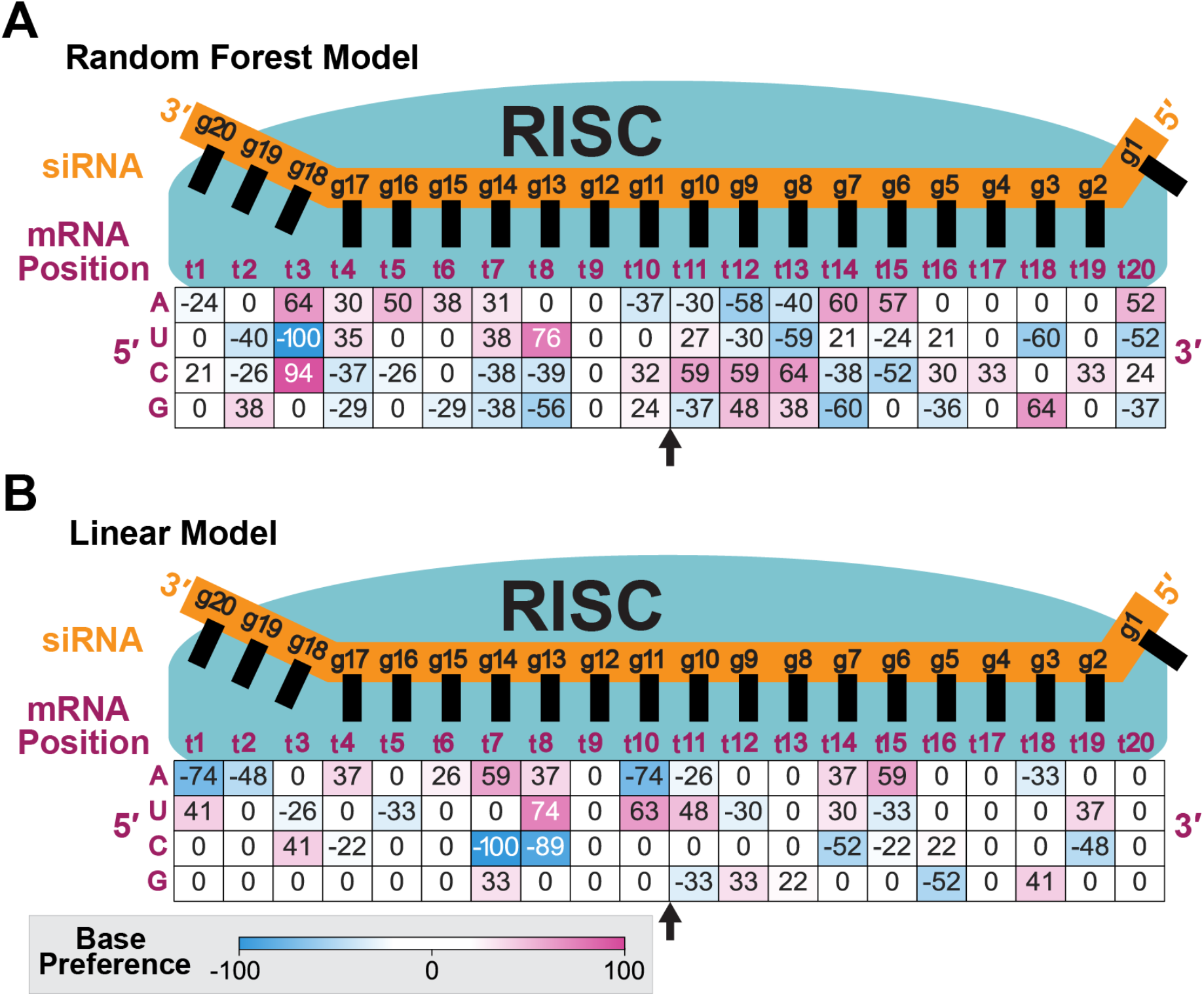
Target site base preferences identified by siRNA efficacy prediction models. (A) Base preferences extracted from random forest model. Positions indicated for target (t) and guide (g) sequences. Larger values indicate greater preference for a base at a position and are colored following the scale indicated. Model developed using 22% effective and 53% ineffective thresholds respectively. Arrow indicates mRNA cleavage site between t10 and t11. (B) Same as A but for a linear model. Base preferences were extracted by proxy (see Results and Methods).

### Comparing base preferences identified by RF vs linear classification models

We then analyzed the extracted base preference matrix from linear and RF models built using the same high-performing threshold pair (Figure 8). Overall trends in base preferences were similar, but RF models produced greater resolution (*i*.*e*., the differences between maximal and minimal base preferences are greater, enabling greater discrimination), likely explaining better model performance. For both models, there was a trend of no base preference near the seed (g2-g5/t16-t19), followed by a region of flexibility (g6-g7/t14-t15), then high affinity near the cleavage site (g8-g11/t10-t13), and high flexibility in the tail (g13-g17/t4-t8). Weaker base importance in the 3′ region (positions t16-t19), which corresponds to the 5′ end of the siRNA seed, likely reflects the need for sequence variability to accommodate a wide range of siRNA, as this region determines siRNA specificity. The lack of specificity here also highlights the flexibility of the model to accommodate a large range of mRNA targets. There is no preference for any base at position t9. Base pairing in this central region is known to be important for effective cleavage (23, 24, 40), therefore it is possible that the lack of preference at this position is necessary to accommodate different bases in different siRNAs and targets. Even a linear model developed using data from unmodified siRNAs showed low preference for any base at position 9 (4), suggesting the importance of this position.

To examine thermodynamic trends, the summed GC preferences were subtracted from the summed AU preferences at each position (Figure 9). At all but three positions (t5, t11, t19) the AU−GC subtracted base preferences had the same directionality in both models. In positions of identical directionality, RF frequently had a larger magnitude. Although thermodynamic asymmetry is required for siRNA efficacy (12), asymmetry was introduced through chemical modification and structure in this siRNA dataset; and thus, does not appear in the weight matrix as a major determinate of efficacy.

**Figure 9.**
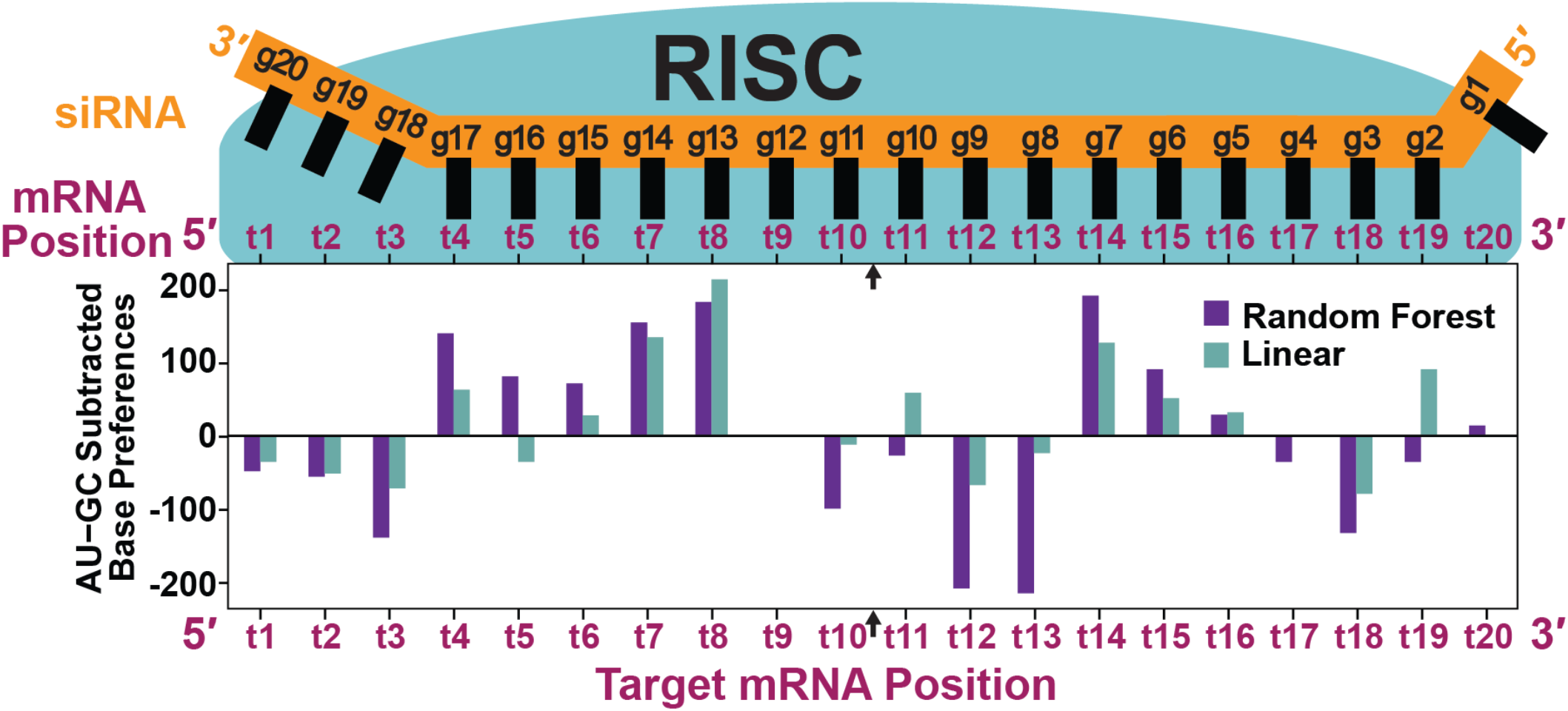
Thermodynamic trends in base preferences extracted from random forest and linear models. Comparing thermodynamic trends approximated by subtracting summed GC preferences from summed AU preferences extracted from the random forest (purple) and linear (teal) models. Positions in sequences indicated for mRNA target (t) and siRNA guide (g) strands. Base preferences were extracted from their respective models using proxy method (see Results and Methods). Base preference positions are indicated with respect to the target mRNA sequence (x-axis). Both linear and random forest models from which preferences were derived were developed using an effective threshold of 22% and an ineffective threshold of 53%.

Overall, base preferences from both models are consistent with the current understanding of siRNA-RISC targeting recognition and cleavage (40, 41).

## DISCUSSION

In this paper, we provide a framework for simple application of ML to small biological datasets. This framework uses a noncanonical trichotomous partitioning method that explores a range of classification thresholds to overcome data variability, uncertainty, and noise common to small biological datasets, a K-fold cross-validation method for training ML models to small datasets that can be adapted (*e*.*g*., changing partition size or K size) to a range of biological problems, and a novel evaluation metric that accounts for data imbalances and varying classification thresholds to enable simple performance comparisons across models. Finally, we present a novel method to extract feature preferences from any classification model by proxy. This framework is presented through the lens of siRNA design but is applicable to any small, variable biological dataset, providing a tool to tap into a previously inaccessible resource to advance biological knowledge.

When using modeling for analysis of complex biological datasets, it is essential to consider the data and the question(s) seeking to be answered in a holistic way. For siRNA design, we selected a classification model type despite siRNA efficacy data being continuous, which is typically better suited for regression. Indeed, many existing siRNA design models apply regression (12, 21, 42, 43). Yet, regression models rely greatly on moderate-efficacy siRNAs, which are not well understood but likely limited by ineffective RISC loading, target release (41, 44, 45), and other complex components that sequence-centered models cannot capture, and cannot be determined from efficacy data alone. Thus, fitting a regression model to these data introduces a large amount of uncertainty into the model, reducing its predictive power. Using classification improves upon existing siRNA prediction models by allowing the model to use data with a clearer underlying mechanism: effective and ineffective siRNAs. In fact, we found that the top-performing model excludes moderate efficacy siRNAs from training.

A major challenge of building a classification model with continuous, inherently noisy siRNA efficacy data is distinguishing data points near a classification threshold. Our trichotomous classification system employs two thresholds – one to define effective siRNA and one to define ineffective – to eliminate noise overlap and enable proper separation of the siRNA classes. By considering a range of effective and ineffective threshold pairs (45 total combinations), we overcome bias associated with selecting a threshold in continuous data not typically amenable to binary classification. We found the most stringent thresholds, which eliminate up to 80% of the training data, provided models with the greatest performance, exemplifying a tradeoff between accuracy and coverage in model building. With a range of thresholds to choose from, we were able to balance satisfactory performance with inclusion of sufficient data to capture biological features driving siRNA efficacy. Trichotomous partitioning also provides greater power over threshold definition, enabling tuning of a model specifically to the problem at hand. For example, by picking a stringent h_1_ threshold and a less stringent h_2_ threshold, the model will more heavily weigh effective siRNAs, enabling a stronger ability to identify them over ineffective siRNAs, which are mostly irrelevant to the biological problem at hand.

Proper model evaluation is a critical step in model building as it identifies a highly predictive model amongst weaker models. Here, AUCPR_adj_ adjusts the commonly used AUCPR by the P_R=1_ to provide a single, easy-to-compare numeric metric that enables performance comparisons across imbalanced datasets and multiple h_1_ thresholds, something other existing model evaluation metrics cannot do. The ability to tailor evaluation metrics to the biological data and question being considered enables proper tuning of a model to optimize its performance. For siRNA design, AUCPR_adj_ prevents overestimation of model performance, and ensures proper model assessment to guide model tuning and final model selection such that the final model performs well when applied in decision making.

We successfully developed a ML model that predicts siRNA efficacy with higher power than a simpler linear model (using the same classification threshold pairs), confirming the power of ML models when applied to biological problems. However, the complexity of ML models makes it challenging to elucidate how they fit the data. This poses a major challenge in cases where the models fit to irrelevant patterns in the data, leading to the development of a poor-quality model that shows high performance (16). Treating such models as “black boxes” limits insight into the biological mechanism the model seeks to describe. The feature extraction method presented here, which extracts proxy features from the model to compute feature preferences, unlocks the potential of using (and examining) more complex models to address important biological questions. Moreover, because our method is evaluation-focused, relying only on the classification outputs of a model, it is simple to apply and adapt to any model type. Having a universal methodology to quickly examine a range of models from the simplest linear models to the most complex deep learning models will shed light into the black box of a model’s predictive mechanism.

Feature preferences from the top performing ML model reflect the current understanding of the RISC mechanism, such as high affinity at the cleavage site, and variability 3′ to the cleavage site (position t9), demonstrating the ability of the model to accurately recapitulate biology. Features extracted from the linear and ML models show similarities in feature preferences, with ML achieving higher resolution.

The linear model is constructed by subtracting base frequencies, and therefore the features will closely reflect the base preferences of the model (Supplementary Figure S8) (4). While the RF model generally aligns with base preferences of the linear model, it also reflects more complex interactions. This is because the linear model considers each base and position entirely independently, whereas the RF model can model feature dependencies through its branching structure (18). In many base-positions where the models differ, the linear model has a weight of zero. This could reflect the linear model’s simplicity (and therefore weakness) in fitting the complex data, leading to washing out of some base-preferences. Moreover, when considering thermodynamic trends derived from feature preferences, both models recapitulate preference for flexibility in the tail that is shown to promote product release (Figure 9) (40, 41). However, RF frequently had larger magnitude trends, which may indicate that the more complex RF model is better at accounting for thermodynamic effects. Our findings suggest the importance of applying ML models to smaller biological datasets as the resulting models can be more biologically accurate and informative (18). While the model itself, not the feature preference matrix, is used for predicting siRNA efficacy, future analysis of feature dependencies and potentially including multiple position-bases in a single feature could further improve the model.

siRNAs are coming of age. Advances in chemical modification have enabled siRNA delivery to many tissues (liver, kidney, brain) (5). A critical next step is identifying effective sequence targets for these siRNA chemistries. Accurate methods to streamline the design and validation of chemically modified siRNAs are needed to complete this task. By applying the framework presented here using well-documented and ready-to-use ML packages like Scikit-Learn (27), or software tools like Weka that require no coding knowledge (46), a wide range of scientists can now harness the power of ML to simplify siRNA drug development.

## SUPPLEMENTARY DATA

Supplementary Data are available at NAR online.

## ACKNOWLEDGEMENT

We thank all Khvorova and Korkin lab members for insightful discussions and support, and Emily Haberlin for helping with the manuscript writing and editing.

