## Supplementary Figures and Tables for "Asymmetric trichotomous data partitioning enables development of predictive machine learning models using limited siRNA efficacy datasets"

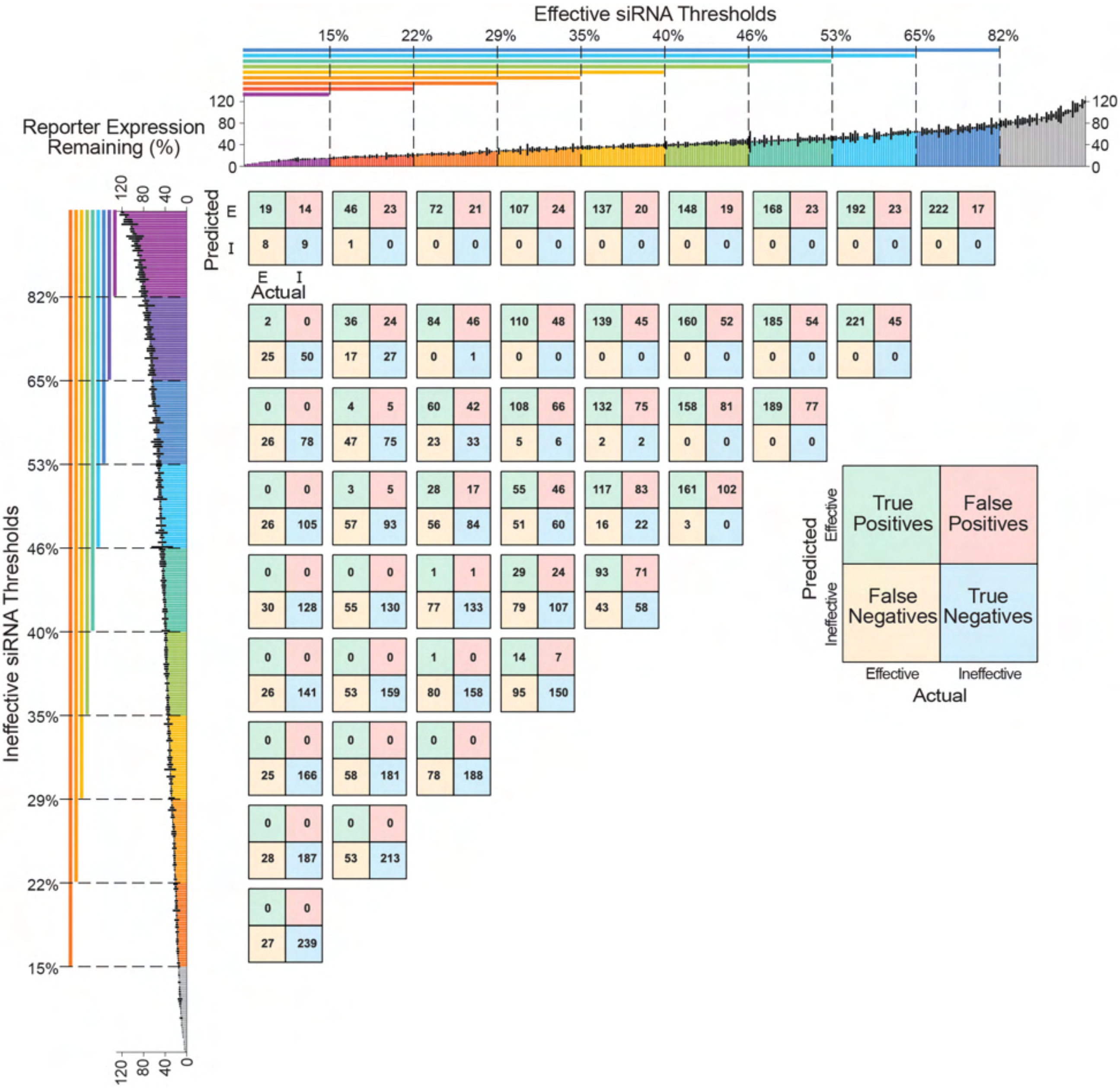


**Supplementary Figure 1. Contingency tables from K-fold cross-validation per classification threshold.** Contingency tables depicting the distribution of actual and random forest classifier-predicted siRNA efficacies. Each table represents a single random forest classifier trained with different effective and ineffective siRNA threshold combinations evaluated on the $K^{th}$ test set. Evaluations on each $K^{th}$ subset were averaged over all K (K=10) rounds of cross-validation. Matrices are color-coded to depict classification group type as indicated in the example larger matrix on the right. Bar plots at top and left depict all siRNA target expression data (as in Figure 2D) colored by effective (top) or ineffective (left) thresholds. Tables are aligned to these bar plots to indicate the effective and ineffective thresholds used for training of that curve's classifier. Thresholds are inclusive of all data with expression values less than (for effective thresholds) or greater than (for ineffective thresholds) the threshold expression percentage. Grey bars indicate siRNAs excluded from model training for the indicated classification (effective or ineffective). Contingency tables were built at the 0.5 probability threshold for all models.

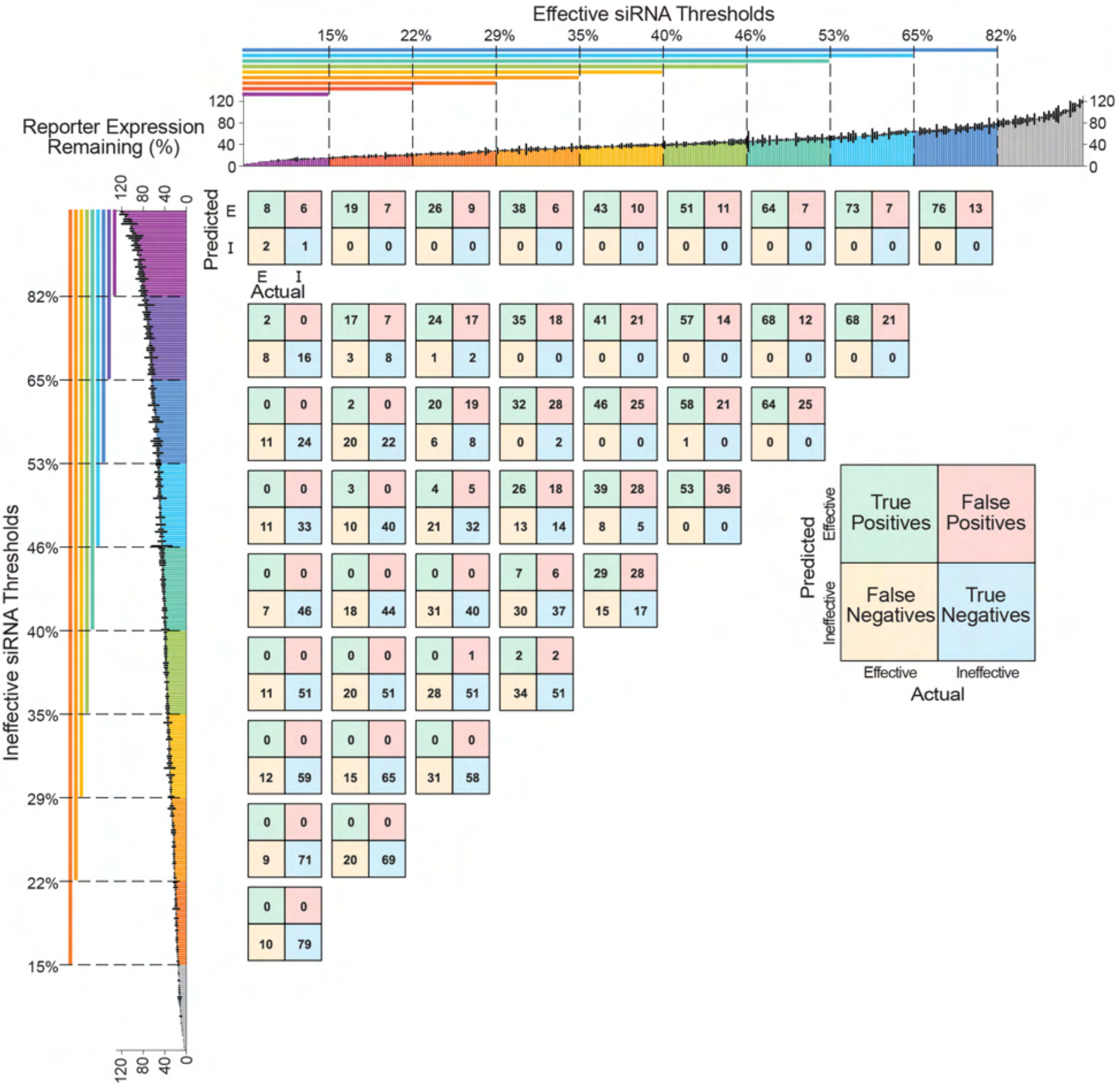


**Supplementary Figure 2. Contingency tables per classification threshold.** Contingency tables depict the actual (bottom) and random forest classifier-predicted (left) distribution of effective and ineffective siRNAs. Each matrix represents a single random forest classifier trained with different effective and ineffective siRNA threshold combinations evaluated on the holdout dataset. Matrices are color-coded to depict classification group type as indicated in the example larger matrix on the right. Bar plots at top and left depict all siRNA target expression data (as in Figure 2D) colored by effective (top) or ineffective (left) thresholds. Tables are aligned to these bar plots to indicate the effective and ineffective thresholds used for training of that curve's classifier. Thresholds are inclusive of all data with expression values less than (for effective thresholds) or greater than (for ineffective thresholds) the threshold expression percentage. Grey bars indicate siRNAs excluded from model training for the indicated classification (effective or ineffective). Contingency tables were built at the 0.5 probability threshold for all models.

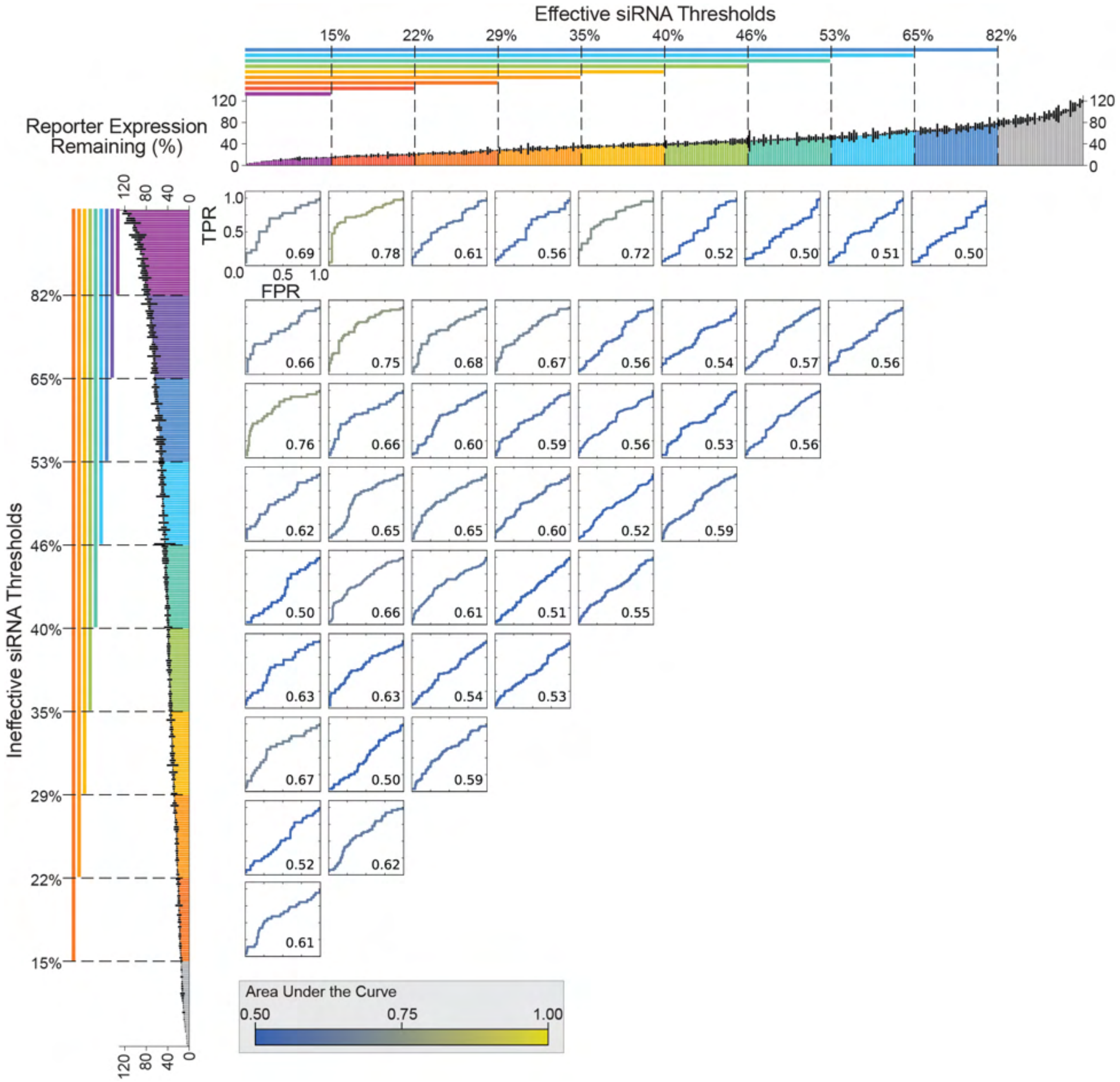


**Supplementary Figure 3. Receiver operating characteristic curves from K-fold cross-validation.** True positive rate (TPR) plotted against false positive rate (FPR) of random forest classifiers evaluated on the K$^{th}$ test set. Evaluations on each K$^{th}$ subset were averaged over all K (K=10) rounds of cross-validation. Each curve represents a single random forest classifier trained with different effective and ineffective siRNA threshold combinations. Curves are colored by the area under the curve. Color bar depicts area under the curve. Bar plots at top and left depict all siRNA target expression data (as in Figure 2D) colored by effective (top) or ineffective (left) thresholds. Curves are aligned to these bar plots to indicate the effective and ineffective thresholds used for training of that curve's classifier. Thresholds are inclusive of all data with expression values less than (for effective thresholds) or greater than (for ineffective thresholds) the threshold expression percentage. Grey bars indicate siRNAs excluded from model training for the indicated classification (effective or ineffective). Area under the curve indicated in bottom right corner of each plot.

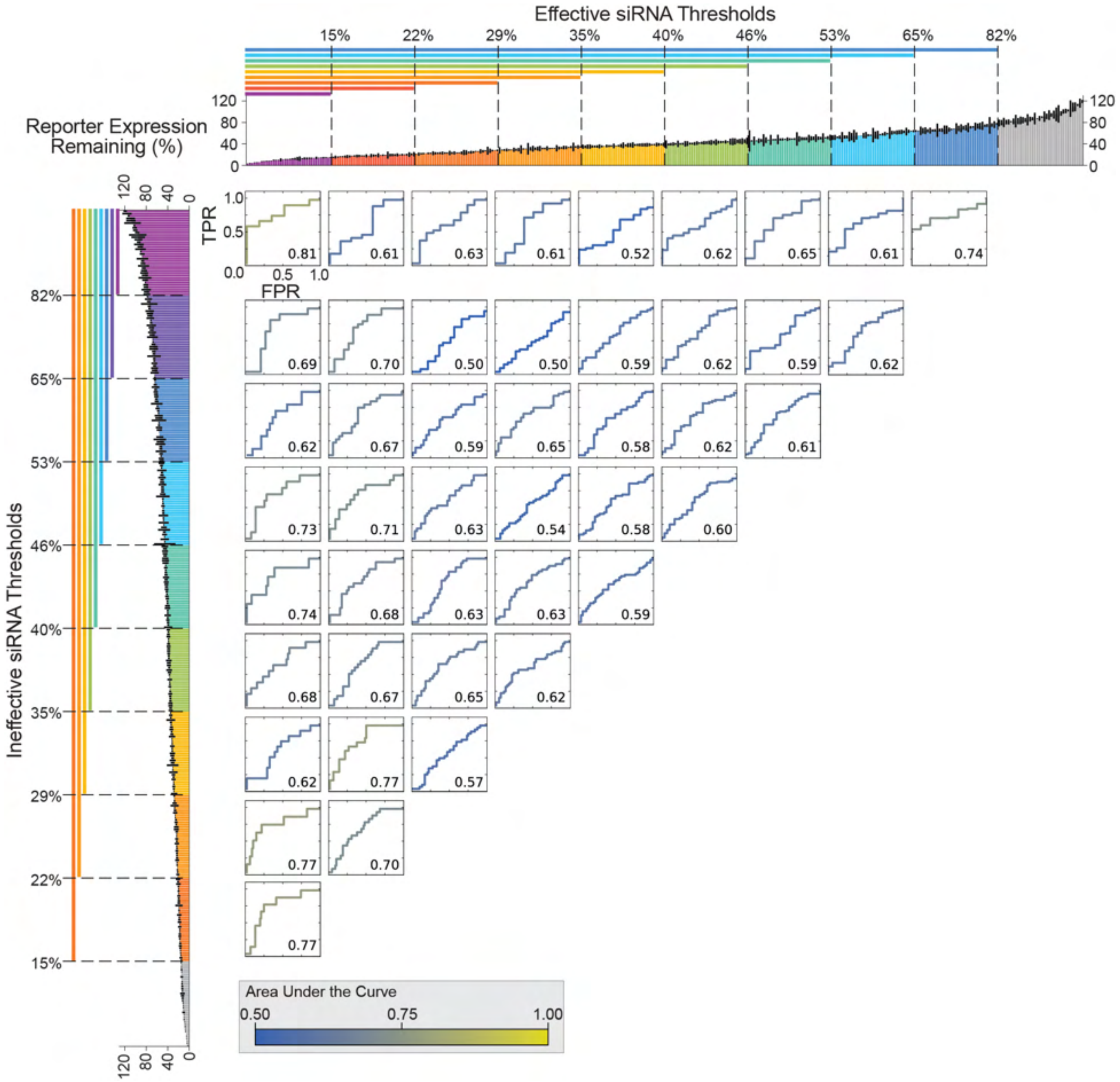


**Supplementary Figure 4. Receiver operating characteristic curves per classification threshold.** True positive rate (TPR) plotted against false positive rate (FPR) of random forest classifiers evaluated on the holdout dataset. Each curve represents a single random forest classifier trained with different effective and ineffective siRNA threshold combinations. Curves are colored by the area under the curve. Color bar depicts area under the curve. Bar plots at top and left depict all siRNA target expression data (as in Figure 2D) colored by effective (top) or ineffective (left) thresholds. Curves are aligned to these bar plots to indicate the effective and ineffective thresholds used for training of that curve's classifier. Thresholds are inclusive of all data with expression values less than (for effective thresholds) or greater than (for ineffective thresholds) the threshold expression percentage. Grey bars indicate siRNAs excluded from model training for the indicated classification (effective or ineffective). Area under the curve indicated in bottom right corner of each plot.

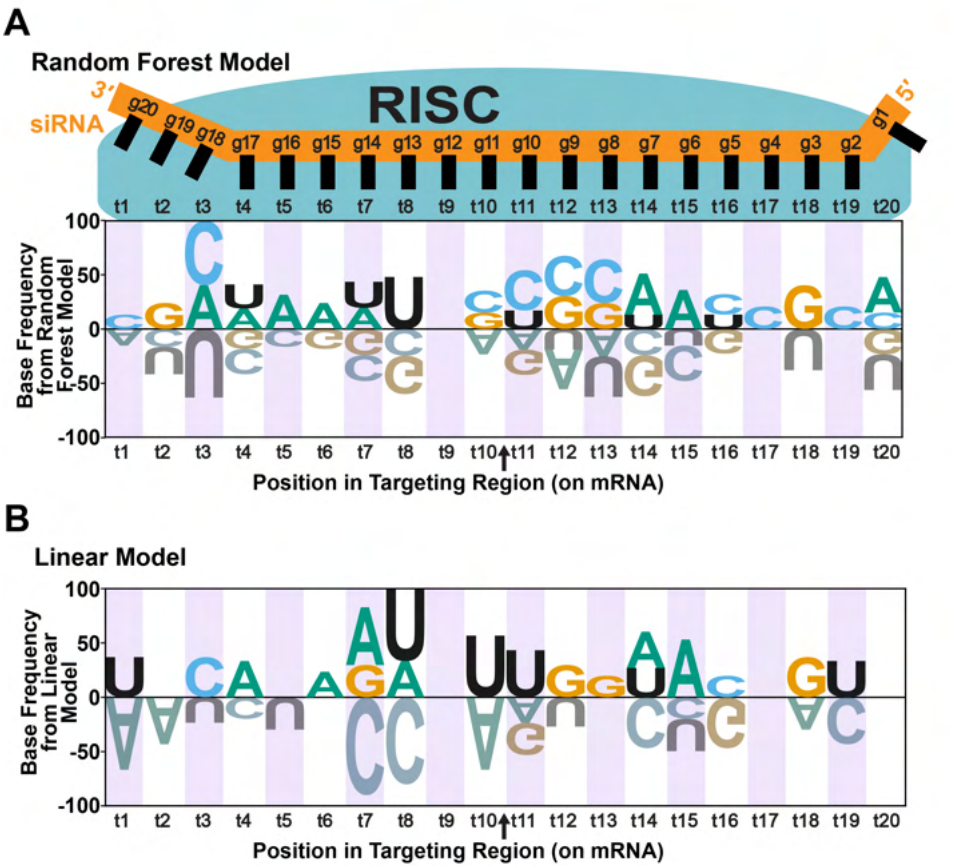


**Supplementary Figure 5. Target site base frequencies identified by siRNA efficacy prediction models depicted as sequence logos.** Base preferences extracted from (**A**) random forest machine learning model or (**B**) linear model using proxy base extraction method (see Results and Methods). Positions in sequences indicated for mRNA target (t) and siRNA guide (g) strands. Letter heights indicate magnitude and direction of base preference at each position; letters oriented upward indicate a favored base, letters oriented downward indicate a disfavored base. Both random forest and linear models were developed using 22% effective and 53% ineffective thresholds respectively. Arrow indicates mRNA cleavage site between t10 and t11. Data depicted here are identical to those presented in Figure 8, but in sequence logo form rather than matrix form.

**A** Base Preferences Extracted Using Proxy Method

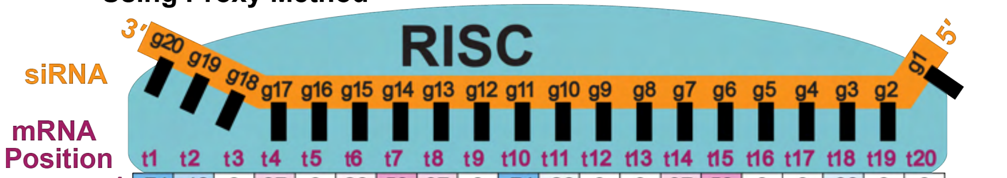


| 5′ | t1 | t2 | t3 | t4 | t5 | t6 | t7 | t8 | t9 | t10 | t11 | t12 | t13 | t14 | t15 | t16 | t17 | t18 | t19 | t20 | 3′ |
|---|---|---|---|---|---|---|---|---|---|---|---|---|---|---|---|---|---|---|---|---|---|
| A | -74 | -48 | 0 | 37 | 0 | 26 | 59 | 37 | 0 | -74 | -26 | 0 | 0 | 37 | 59 | 0 | 0 | -33 | 0 | 0 | |
| U | 41 | 0 | -26 | 0 | -33 | 0 | 0 | 74 | 0 | 63 | 48 | -30 | 0 | 30 | -33 | 0 | 0 | 0 | 37 | 0 | |
| C | 0 | 0 | 41 | -22 | 0 | 0 | -100 | -89 | 0 | 0 | 0 | 0 | 0 | -52 | -22 | 22 | 0 | 0 | -48 | 0 | |
| G | 0 | 0 | 0 | 0 | 0 | 0 | 33 | 0 | 0 | 0 | -33 | 33 | 22 | 0 | 0 | -52 | 0 | 41 | 0 | 0 | |

**B** Base Preferences Derived Directly from Linear Model Coefficeints

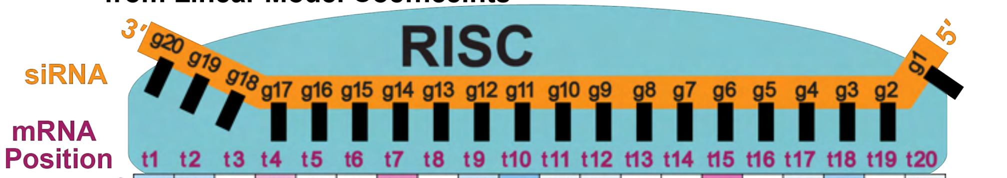


| 5′ | t1 | t2 | t3 | t4 | t5 | t6 | t7 | t8 | t9 | t10 | t11 | t12 | t13 | t14 | t15 | t16 | t17 | t18 | t19 | t20 | 3′ |
|---|---|---|---|---|---|---|---|---|---|---|---|---|---|---|---|---|---|---|---|---|---|
| A | -49 | -45 | 0 | 49 | -24 | 0 | 68 | 0 | -42 | -66 | -29 | -30 | 21 | 0 | 78 | 0 | -31 | -56 | 0 | -28 | |
| U | 91 | 0 | 0 | 0 | 0 | 0 | 0 | 84 | 21 | 67 | 51 | -41 | 0 | 0 | -88 | 33 | 29 | 27 | 0 | 28 | |
| C | -27 | 30 | 0 | -27 | 0 | 0 | -100 | -88 | 23 | 0 | 0 | 38 | 0 | -30 | 0 | 26 | 0 | 0 | 0 | 0 | |
| G | 0 | 0 | 0 | 0 | 23 | 0 | 37 | 0 | 0 | 0 | 0 | 33 | -22 | 0 | 0 | -65 | 0 | 34 | 0 | 0 | |

Base Preference -100 0 100

**Supplementary Figure 6. Proxy method for base preference extraction shows high correlation with preferences directly extracted from linear model.** Base preferences extracted from the same linear model using (**A**) the proxy feature extraction method and (**B**) directly from the linear model coefficients (see Results and Methods). Positions in sequences indicated for mRNA target (t) and siRNA guide (g) strands. Base preferences are color coded following the scale indicated at the bottom; a larger positive value indicates stronger base preference and more negative values indicate greater base disfavoring at a position. Both matrices were derived from the same linear model developed using 22% effective and 53% ineffective thresholds respectively. Arrow indicates mRNA cleavage site between t10 and t11.

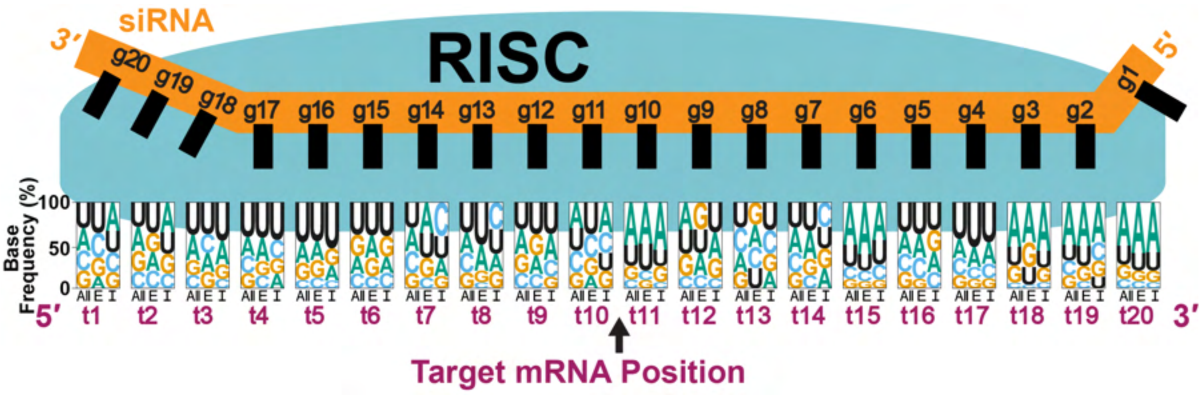


**Supplementary Figure 7. siRNA base frequencies compared across effective and ineffective siRNAs in top performing threshold pair.** Base frequencies for: all 356 siRNAs in the dataset used for analysis (**All**), only those 71 identified as effective by the 22% threshold (**E**; effective), or only those 103 identified as ineffective by the 53% threshold (**I**; ineffective). Positions in sequences indicated for mRNA target (t) and siRNA guide (g) strands. Letter heights and top-to-bottom positioning indicate magnitude base frequency at each position with most frequent base at a position appearing at the top and with the largest height. Arrow indicates mRNA cleavage site between t10 and t11.

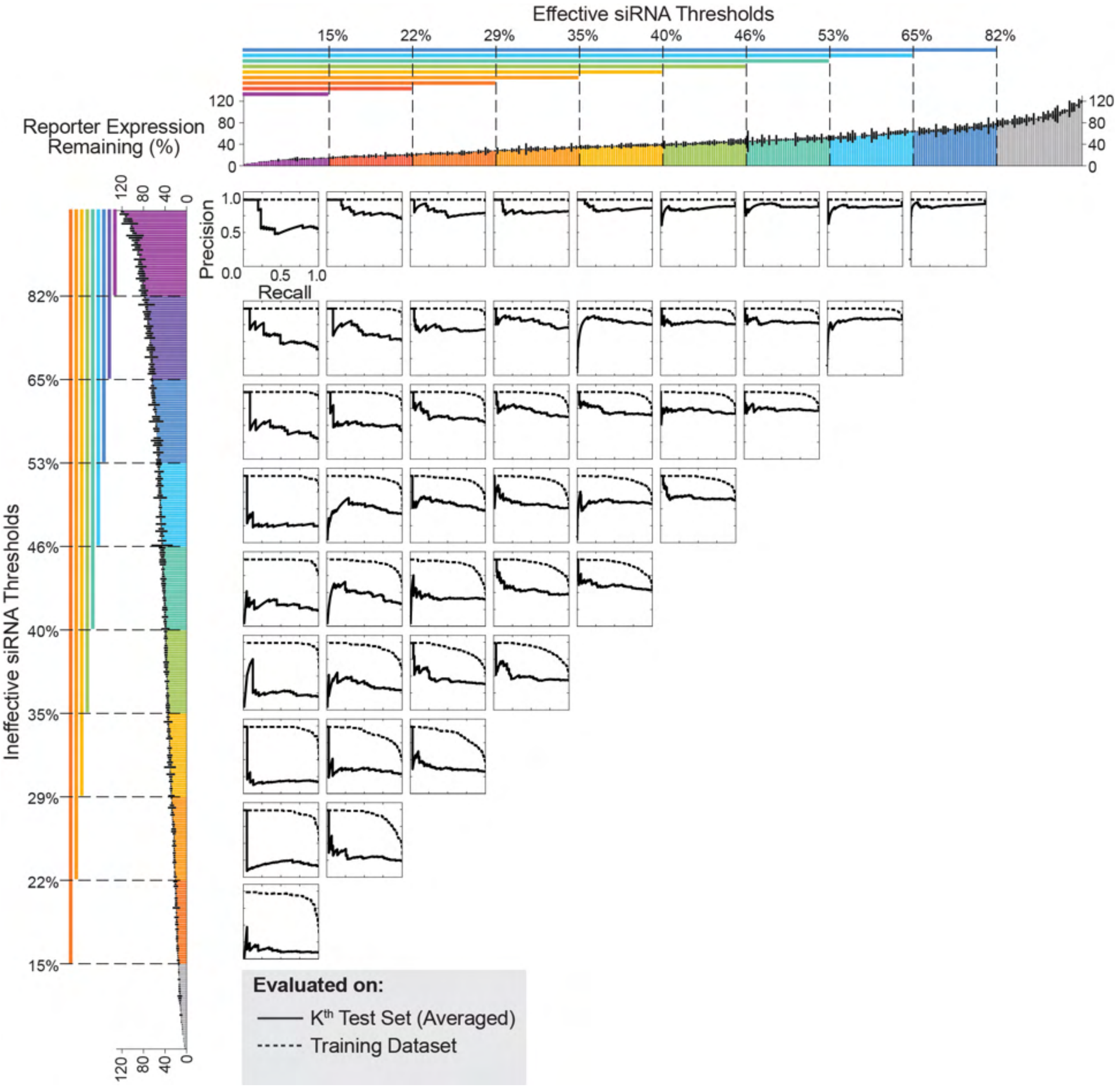


**Supplementary Figure 8. Comparing average model performances on training vs test sets during K-fold cross-validation.** Precision recall curves for random forest classifiers evaluated on the training dataset (dotted curves) and the $K^{th}$ test set (solid curves). Each set of overlaid curves represents a single random forest classifier trained with different effective and ineffective siRNA threshold combinations. Bar plots at top and left depict all siRNA target expression data (as in Figure 2D) colored by effective (top) or ineffective (left) thresholds. Curves are aligned to these bar plots to indicate the effective and ineffective thresholds used for training of that curve's classifier. Thresholds are inclusive of all data with expression values less than (for effective thresholds) or greater than (for ineffective thresholds) the threshold expression percentage. Grey bars indicate siRNAs excluded from model training for the indicated classification (effective or ineffective). Evaluations on each $K^{th}$ subset were averaged over all K (K=10) rounds of cross-validation.

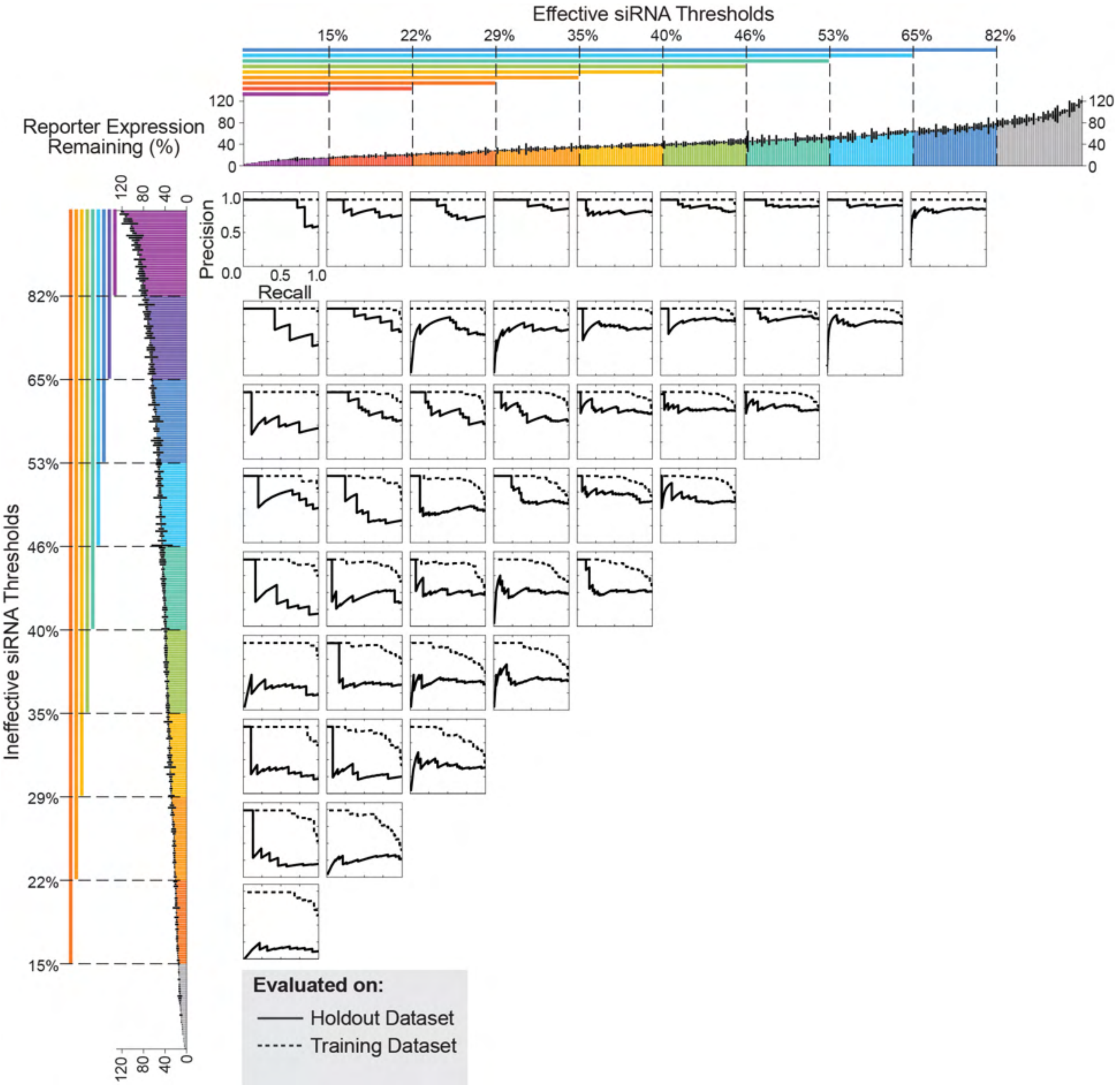


**Supplementary Figure 9. Comparing model performance on holdout dataset vs training dataset per classification threshold.** Precision recall curves for random forest classifiers evaluated on the training dataset (dotted curves) and the holdout dataset (solid curves). Each set of overlaid curves represents a single random forest classifier trained with different effective and ineffective siRNA threshold combinations. Bar plots at top and left depict all siRNA target expression data (as in Figure 2D) colored by effective (top) or ineffective (left) thresholds. Curves are aligned to these bar plots to indicate the effective and ineffective thresholds used for training of that curve's classifier. Thresholds are inclusive of all data with expression values less than (for effective thresholds) or greater than (for ineffective thresholds) the threshold expression percentage. Grey bars indicate siRNAs excluded from model training for the indicated classification (effective or ineffective).

| Normalized $AUCPR_{adj}$ | Precision at Recall=1 ($P_{R=1}$) | AUCPR | Effective Threshold (% Reporter Expression Remaining) | Ineffective Threshold (% Reporter Expression Remaining) |
|---|---|---|---|---|
| 100 | 0.588 | 0.904 | 80 | 15 |
| 99 | 0.435 | 0.75 | 65 | 15 |
| 84 | 0.149 | 0.42 | 40 | 15 |
| 74 | 0.645 | 0.873 | 65 | 22 |
| 72 | 0.302 | 0.53 | 46 | 22 |
| 67 | 0.564 | 0.863 | 52 | 22 |
| 66 | 0.5 | 0.704 | 46 | 15 |
| 64 | 0.51 | 0.708 | 52 | 29 |
| 52 | 0.182 | 0.345 | 29 | 15 |
| 48 | 0.565 | 0.707 | 46 | 35 |
| 42 | 0.552 | 0.672 | 52 | 35 |
| 41 | 0.158 | 0.283 | 22 | 15 |
| 40 | 0.603 | 0.718 | 46 | 40 |
| 39 | 0.327 | 0.442 | 40 | 22 |
| 37 | 0.743 | 0.844 | 80 | 29 |
| 34 | 0.351 | 0.449 | 35 | 22 |
| 34 | 0.595 | 0.691 | 65 | 29 |
| 33 | 0.19 | 0.287 | 35 | 15 |
| 31 | 0.44 | 0.526 | 52 | 15 |
| 31 | 0.823 | 0.905 | 80 | 46 |
| 28 | 0.76 | 0.832 | 80 | 22 |
| 28 | 0.217 | 0.297 | 29 | 22 |
| 28 | 0.449 | 0.526 | 40 | 29 |
| 28 | 0.864 | 0.933 | 80 | 35 |
| 28 | 0.667 | 0.74 | 52 | 40 |
| 27 | 0.5 | 0.571 | 40 | 40 |
| 24 | 0.455 | 0.519 | 46 | 29 |
| 22 | 0.596 | 0.65 | 46 | 46 |
| 22 | 0.719 | 0.773 | 52 | 52 |
| 17 | 0.738 | 0.774 | 52 | 46 |
| 16 | 0.354 | 0.39 | 35 | 29 |
| 16 | 0.901 | 0.929 | 80 | 52 |
| 16 | 0.85 | 0.881 | 65 | 52 |
| 15 | 0.695 | 0.724 | 65 | 40 |
| 14 | 0.119 | 0.151 | 15 | 15 |
| 14 | 0.463 | 0.492 | 40 | 35 |
| 14 | 0.913 | 0.933 | 80 | 65 |
| 13 | 0.36 | 0.388 | 29 | 29 |
| 13 | 0.803 | 0.824 | 65 | 46 |
| 12 | 0.409 | 0.432 | 35 | 35 |
| 11 | 0.811 | 0.825 | 80 | 40 |
| 9 | 0.764 | 0.773 | 65 | 65 |
| 3 | 0.673 | 0.66 | 65 | 35 |
| 1 | 0.241 | 0.23 | 22 | 22 |
| 0 | 0.854 | 0.828 | 80 | 80 |

**Supplementary Table S1. Normalized adjusted AUCPR, AUCPR, and Precision at Recall = 1 for each threshold pair evaluated on holdout set.** AUCPR and precision at recall=1 were used to compute the normalized $AUCPR_{adj}$. Each row represents a threshold pair that is defined by the stated effective threshold (2nd to last column) and effective threshold (last column).

| Normalized $AUCPR_{adj}$ | Precision at Recall=1 ($P_{R=1}$) | AUCPR | Effective Threshold (% Reporter Expression Remaining) | Ineffective Threshold (% Reporter Expression Remaining) |
|---|---|---|---|---|
| 100 | 0.365 | 0.573 | 15 | 65 |
| 83 | 0.277 | 0.434 | 15 | 52 |
| 73 | 0.515 | 0.625 | 22 | 65 |
| 72 | 0.701 | 0.828 | 22 | 80 |
| 64 | 0.306 | 0.447 | 22 | 40 |
| 57 | 0.395 | 0.563 | 22 | 52 |
| 57 | 0.525 | 0.636 | 29 | 52 |
| 57 | 0.457 | 0.572 | 29 | 46 |
| 57 | 0.696 | 0.794 | 35 | 65 |
| 54 | 0.617 | 0.715 | 35 | 52 |
| 48 | 0.551 | 0.64 | 15 | 80 |
| 46 | 0.36 | 0.461 | 29 | 35 |
| 43 | 0.638 | 0.712 | 40 | 52 |
| 42 | 0.454 | 0.539 | 35 | 40 |
| 40 | 0.403 | 0.489 | 22 | 46 |
| 38 | 0.521 | 0.593 | 40 | 40 |
| 37 | 0.502 | 0.574 | 35 | 46 |
| 37 | 0.413 | 0.492 | 35 | 35 |
| 35 | 0.262 | 0.348 | 22 | 35 |
| 33 | 0.205 | 0.291 | 22 | 22 |
| 33 | 0.617 | 0.673 | 46 | 46 |
| 32 | 0.299 | 0.375 | 29 | 29 |
| 30 | 0.2 | 0.28 | 15 | 40 |
| 28 | 0.755 | 0.789 | 40 | 65 |
| 28 | 0.664 | 0.706 | 46 | 52 |
| 27 | 0.162 | 0.24 | 15 | 35 |
| 24 | 0.261 | 0.325 | 22 | 29 |
| 24 | 0.758 | 0.784 | 46 | 65 |
| 22 | 0.798 | 0.817 | 29 | 80 |
| 22 | 0.774 | 0.796 | 52 | 65 |
| 21 | 0.711 | 0.735 | 52 | 52 |
| 20 | 0.892 | 0.9 | 52 | 80 |
| 18 | 0.125 | 0.188 | 15 | 22 |
| 17 | 0.823 | 0.83 | 35 | 80 |
| 16 | 0.677 | 0.694 | 29 | 65 |
| 14 | 0.215 | 0.263 | 15 | 46 |
| 14 | 0.873 | 0.871 | 40 | 80 |
| 14 | 0.559 | 0.581 | 40 | 46 |
| 12 | 0.373 | 0.405 | 29 | 40 |
| 7 | 0.135 | 0.174 | 15 | 29 |
| 7 | 0.904 | 0.884 | 65 | 80 |
| 3 | 0.831 | 0.81 | 65 | 65 |
| 2 | 0.104 | 0.136 | 15 | 15 |
| 2 | 0.936 | 0.904 | 80 | 80 |
| 0 | 0.897 | 0.864 | 46 | 80 |

**Supplementary Table S2. Normalized adjusted AUCPR, AUCPR, and Precision at Recall = 1 for each threshold from K-fold cross-validation.** AUCPR and precision at recall=1 were used to compute the normalized $AUPCR_{adj}$. Each row represents a threshold pair that is defined by the stated effective threshold (2nd to last column) and effective threshold (last column).
